# Transcriptional dysregulation and impaired neuronal activity in *FMR1* knock-out and Fragile X patients’ iPSC-derived models

**DOI:** 10.1101/2023.08.30.554628

**Authors:** Gilles Maussion, Cecilia Rocha, Narges Abdian, Dimitri Yang, Julien Turk, Dulce Carrillo Valenzuela, Luisa Pimentel, Zhipeng You, Barbara Morquette, Michael Nicouleau, Eric Deneault, Samuel Higgins, Carol X.-Q. Chen, Wolfgang Reintsch, Ho Stanley, Vincent Soubannier, Sarah Lépine, Zora Modrusan, Jessica Lund, William Stephenson, Rajib Schubert, Thomas M. Durcan

## Abstract

The lack of fragile X mental retardation protein (FMRP) protein, due to a repression of the *FMR1* gene, causes Fragile X syndrome (FXS), one of the most prevalent forms of syndromic autisms. The *FMR1* gene codes for an RNA binding protein involved in the regulation of gene expression through RNA processing, control of local translation, and protein-protein interactions; processes that are crucial for proper brain development.

Taking advantage of induced pluripotent stem cells (iPSCs) and CRISPR-Cas9 genome editing technologies, we generated iPSC-derived cortical neural progenitors and cortical neurons from an *FMR1* knock-out and patient cell line with the aim of identifying common phenotypes between the two cellular models. Using RNA sequencing, quantitative PCR and multielectrode array approaches, we assessed how the absence of the functional *FMR1* gene affects the transcriptional profiles and the activities of iPSC-derived cortical neuronal progenitor cells (NPCs) and neurons with both models.

We observed that *FMR1* KO and FXS patient cells have a decrease in their mean firing rate; a cellular activity that can also be blocked by tetrodotoxin (TTX) application in wild-type active neurons. Relative to wild-type neurons, in *FMR1* KO neurons, increased expression of presynaptic mRNA and transcription factors involved in the forebrain specification and decreased levels of mRNA coding AMPA and NMDA subunits were observed. Intriguingly, 40% of the differentially expressed genes were commonly deregulated between NPCs and differentiating neurons with significant enrichments in FMRP targets and Autism Related Genes found amongst downregulated genes. This implies that an absence of functional FMRP affects transcriptional profiles at the NPC stage, resulting in impaired activity and differentiation of the progenitors into mature neurons over time.

These findings from the *FMR1* KO lines were also shared with FXS patients’ iPSC-derived cells that also present with an impairment in activity and neuronal differentiation, illustrating the critical role of FMRP protein in neuronal development.

## Introduction

Fragile X syndrome (FXS) is classified as a syndromic autism and falls in the category of 15 to 20% of autism spectrum disorders (ASDs) whose genetic causes are known [1]. In the majority of cases, FXS is caused by a full repression of the *FMR1* gene expression due to hypermethylation of an extended CGG repeated sequence located in the *5’UTR* region of the gene [2]. The *FMR1* gene is located on the chromosomal region Xq27.3 and encodes the Fragile X Messenger Ribonucleoprotein (FMRP), a multifunctional protein [3–5] that plays a key role in regulating the local translation of proteins required for proper neuronal development. With an estimated prevalence of 1/4000 males and 1/7000 females [6, 7], FXS is the leading cause of intellectual disability worldwide. Patients diagnosed with FXS also exhibit social impairment, developmental delay, hyperactivity, and macroorchidism [8].

Studying the cellular and molecular mechanisms underlying neurodevelopmental disorders is still challenging, but progress is being made as new models to capture early events of brain formation and development become available. Post-mortem brain studies and mouse models, which include both genetic and chemically induced ASD models, have contributed to understanding the pathogenesis of neurodevelopmental disorders, and in particular syndromic forms of ASDs [9]. These studies showed how defects in mitochondrial function, dendritic trafficking, and synaptic connectivity are causative factors in impaired development[10]. However, syndromic forms of autism with known genetics account for only ∼5-15% of the cases [1] highlighting the limitations of the above-mentioned models.

Since the advent of methodologies to reprogram somatic cells into induced pluripotent stem cells (iPSCs) [11], we can now make human neurons and other brain-related cell types in a dish from individual patients, providing us with the tools and patient-derived cell types to investigate the molecular and cellular underpinnings of neurodevelopmental disorders [9, 12]. One major advantage of iPSCs is their potential for personalized medicine as studies can be conducted directly on patient cells ensuring that future therapies can be directed to their own genetic background [13].

Although most studies related to autism spectrum disorders have focused on the properties of iPSC-derived neurons, many also highlighted significant alterations at the neural progenitor stage [14–16]. In a previous study, haploinsufficiency in the *GRIN2B* gene (associated with ASD) leads to a disruption of cortical differentiation, although we also observed GRIN2B expression and NMDA response at the NPC stage [17]. This implies that the NMDA receptor is required at very early stages of neuronal development. Interestingly, altered GRIN2B expression was observed in cortical neurons of an *Fmr1-*deficient mouse model [18]. Hypothesizing that the FMRP protein plays a role in the neuronal development from the neural progenitor stage, we chose to investigate its gene expression and profile the differentiation of progenitors into neurons to better understand how an absence of FMRP impacts cortical neurogenesis with these iPSC-derived cellular model.

Different studies have reported findings with iPSC-derived neurons used to model FXS pathophysiology either from *FMR1* KO lines or from FXS patient lines [19]. However, the increase in the number of iPSC models tested has generated heterogeneous results across studies. These differences can be explained by inter-patient variations such as genetic background, or by experimental procedures, such as cell reprogramming or the differentiation protocol used to generate the cell type of interest [20]. One of the issues that has been raised regarding the investigation of iPSC-derived cortical neurons is the ability to mature those cells, which seems more time-consuming and challenging than other iPSC-derived neuronal subtypes. Issues to successfully obtain mature and electrically active cortical neurons have been reported [21]. Thus, we first compared two differentiation protocols to obtain cortical neurons that could then be tested for their electrical activity.

To identify the phenotypes expressed due to the absence of *FMR1* gene expression, we performed our experiments with an isogenic control, generating a knockout of *FMR1* in a control cell line that was previously fully characterized [22]. Next, neuronal electrical activity was tested, and gene expression profiles were assessed in iPSC-derived cells from *FMR1* KO and FXS patient lines. Taken together, analyses performed on neurons from *FMR1* KO and FXS patient uncovered altered molecular and cellular phenotypes responsible for the neurodevelopmental phenotypes observed in Fragile X syndrome.

## Material and Methods

### Cell line description

The use of the following IPSCs in this research is approved by the McGill University Health Centre Research Ethics Board (DURCAN_IPSC/2019-5374).

**Table 1:** Description of the iPSC lines.

| Cell line | Diagnostic | Cell of origin | Sex | Ethnicity | Reprogramming | Cell Source | References |
| --- | --- | --- | --- | --- | --- | --- | --- |
| AIW002-02 | Healthy | PBMC | M | Caucasian | Episomal | The Neuro | PMID: 34287353 |
| *FMR1*-KO | NA | AiW002-2 IPSC | M | Caucasian | N/A | The Neuro |  |
| FX-11-7 | Fragile X Syndrome | Fibroblast | M | N/A | Lentivirus | WiCell | PMID: 24654675 |

### Cell culture and Cortical neuron differentiation

iPSCs were cultivated in 10 cm dishes pre-coated with Matrigel (Corning) and were maintained in mTeSR1 media (STEMCELL Technologies) with daily media change. Cells were passaged with Gentle Cell Dissociation Reagent (STEMCELL Technologies). Cortical progenitors (NPCs) were obtained from iPSCs as described by [23] and then banked. Progenitor cells were grown in T75 flasks pre-coated with Poly-L-Ornithine (PO) and laminin in NPC progenitor media composed by DMEM-F12 supplemented with N2, B27, NEAA, Antibiotic-Antimycotic, laminin (1 µg/mL), EGF (20 ng/mL) and FGFb (20 ng/mL). For cortical neuronal differentiation, cells received Neurobasal media supplemented with N2, B27, Compound E (0.1 µM), db-cAMP (500 µM), Ascorbic Acid (200 µM), BDNF (20 ng/mL), GDNF (20 ng/mL), TGF-b3 (1 ng/mL), laminin (1 µg/mL) or Forebrain media (STEMCELL Technologies), as summarized in **Figure 1**. Briefly, cells were kept in the appropriate media for four weeks, and half of the medium was changed twice a week. Neurons differentiated with forebrain media were kept in differentiation media-STEMdiff forebrain neuron differentiation medium + supplements for one week and subsequently in maturation media – Brainphys Neuronal Medium + supplements (STEMCELL technologies) for the remaining days in culture, Half of the medium was changed twice a week.

**Figure 1:**
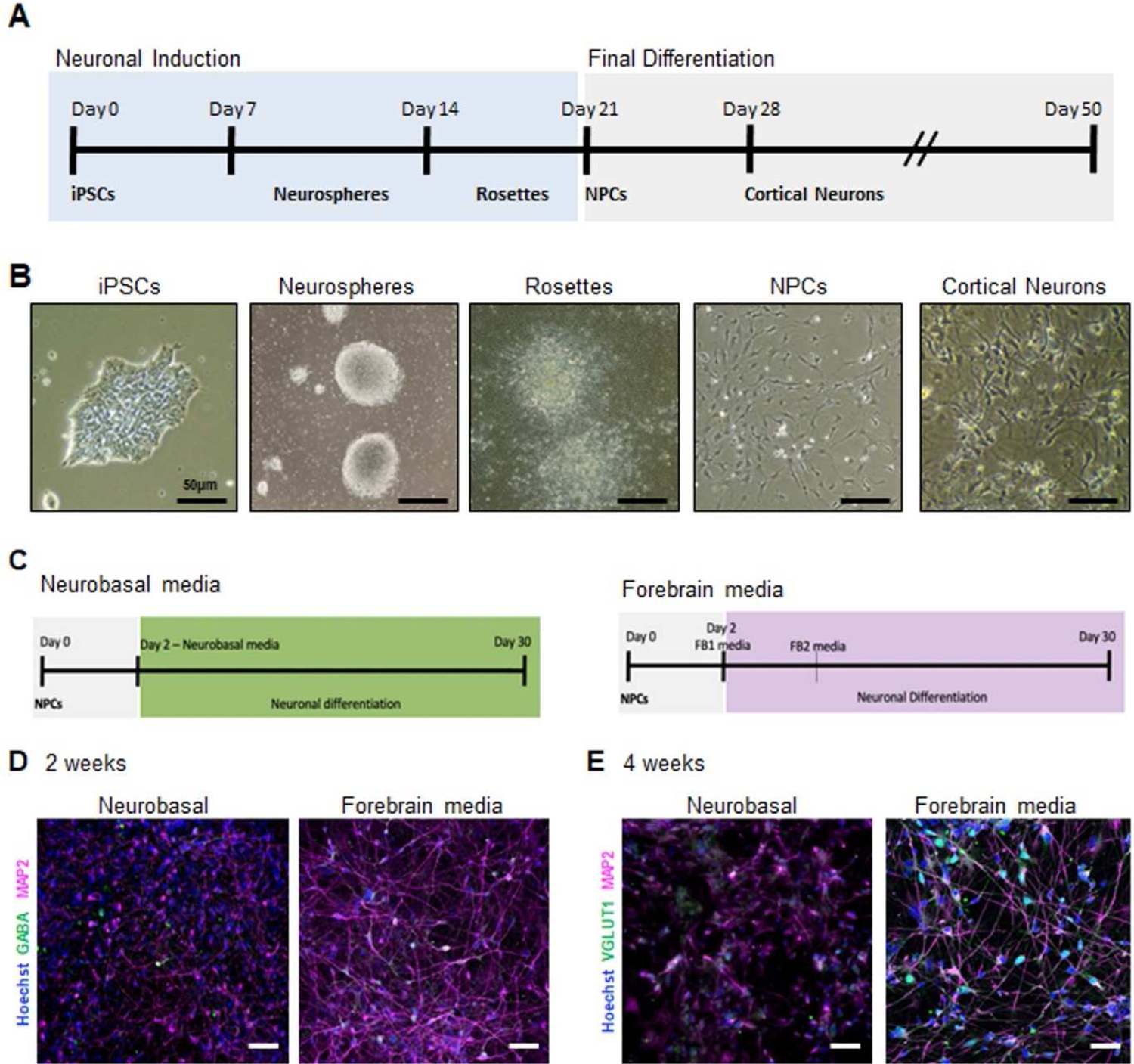
Generation of electrically active iPSC-derived cortical neurons. (A) Schematic of the protocol for the generation of cortical neurons from human iPSCs. (B) Light microscopy images showing the various stages of the generation of cortical neuron generation. Neurospheres are obtained from iPSCs and are then plated forming rosettes that will expand giving rise to NPCs. (C) Schematic of the protocol for the generation of cortical neurons using Neurobasal based and Forebrain media. Immunofluorescence images of (D) 2 weeks and (E) 4 weeks cortical neurons showing expression of the neuronal markers GABA or VGLUT1 (green) and MAP2 (magenta). Nuclei were counterstained with Hoechst. The scale bars are 50 µm long.

### CRISPR genome editing

Synthetic gRNA (sgRNA) was designed using benchling.com [24] to generate one double-strand break (DSB) in the *FMR1* gene (ENSG00000102081) by Cas9 nuclease. DSB is followed by homology-directed repair (HDR) and stop tag insertion (**Supplementary Table 1**). CRISPR/Cas9 editing was achieved when CAS9 protein (1 μl; stock 61 μM; Alt-R® S.p. HiFi Cas9 Nuclease V3, IDT), *FMR1* sgRNA (3 μl; stock 100 μM, Synthego), and the ssODN (1 μl; stock 100 μM, IDT) in 20 μl of nucleofection buffer P3 (P3 Primary Cell NucleofectorTM Solution, Lonza) were nucleofected (program CA137, 4D-Nucleofector Device, Lonza) into 500,000 detached iPSCs [25]. Following nucleofection, iPSCs were evenly distributed into a flat-bottom 96-well plate in mTeSR media and 10 μM Y-27632. After limiting dilution, gene-edited clones were identified by ddPCR (QX200™ Droplet Reader, Bio-Rad) [25] and Sanger sequencing. For more details, see our CRISPR editing [26] and DNA sequencing [27] protocols.

### Immunofluorescence staining and image acquisition

Cells were seeded onto glass coverslips pre-coated with Matrigel (iPSCs) or PO and laminin (NPCs and neurons). Cultures were fixed with 4% paraformaldehyde (PFA) in phosphate-buffered saline (PBS) for 10 minutes at room temperature. Coverslips were then incubated with a blocking solution containing 5% normal donkey serum (NDS), 0.05% bovine serum albumin (BSA), and 0.2% Triton in PBS for 1 hour at room temperature or overnight at 4°C in a humidified chamber. The solution of primary antibodies (**Supplementary Table 2**) was prepared in the blocking solution. Coverslips were then incubated in primary antibody solution overnight at 4°C in a humid chamber. The next day, samples were washed three times for 10 minutes with PBS and incubated with a solution of Alexa Fluor-conjugated secondary antibodies (1:1000 Life Technologies), counterstained with Hoechst (Thermo Fisher Scientific H3570), then mounted in antifade mounting media Aqua-Poly/mount (Polyscience). Images were collected with an EVOS imaging system (Invitrogen EVOS FL Auto 2) and a confocal laser-scanning microscope (Leica SP8). Images were processed and analyzed using ImageJ software. Neuronal networks were assessed by the Integrated density representing the sum of the values of the pixels in each image.

### Multielectrode Array (MEA) activity analysis

Spontaneous neuronal electrical activity was measured by MEA. Cortical neurons and cortical spheres were differentiated in 24 well plates (CytoView MEA 24; Axion) for MEA system. For 2D cultures, 80,000 NPCs were seeded onto PO/laminin pre-coated MEA plates. Cultures were treated with 1 µm Tetrodotoxin (TTX, Abcam) for 30 min. Measurements started a week after plating, and recordings of spontaneous activity were performed once a week for 4 weeks. Different measurements (number of spikes and mean firing rate) were taken during a 5-minute consecutive duration recording on different days using Maestro Edge MEA System (Axion Biosystems). Measurements were analyzed with AxIS Navigator and NeuroExplorer 5 software. Data was then exported to Microsoft Excel software and graphed using the GraphPad Prism software.

### RNA extraction, cDNA synthesis, and quantitative PCR

NPCs and iPSC-derived neurons were dissociated using Accutase^®^ Cell Dissociation Reagent (Thermo Fisher Scientific) incubated at 37°C for 5 minutes. Cells were then collected and harvested by centrifugation for 5 min at 1500 rpm. Cell pellets were resuspended in Qiazol (Qiagen) and stored at − 80 °C before total RNA extraction with the miRNAeasy (Qiagen) kit.

Reverse transcription reactions were performed on 500 ng of total RNA extract to obtain cDNA in a 20 μl total volume containing, using the iScript Reverse Transcription Supermix (Biorad). The reactions were conducted in single plex, in a 10 µl total volume containing 2X TaqMan Fast Advanced Master Mix, 20X TaqMan primers/probe set (Thermo Fisher Scientific), 1 µL of diluted cDNA and RNAse-free H_2_O. Real-time PCR (RT-PCR) were performed on a QuantStudio 3 or QuantStudio 5 machines (Thermo Fisher Scientific). Primers/probe sets from Applied Biosystems were selected from the Thermo Fisher Scientific website (**Supplementary Table 3**). Two endogenous controls (beta-actin and GAPDH) were used for normalization. The normalized expression levels were determined according to the ΔCT method [28]. Data was analyzed with GraphPad Prism. Statistics were processed using either a one-way or two-way ANOVA with post-hoc tests.

### RNA sequencing

#### Sample preparation and sequencing

The cells were dissociated using accutase and lysed in the QIAzol buffer. RNA was extracted using the miRNeasy from QIAGEN and quantified using NanoDrop (ThermoFisher Scientific). Their quality was assessed using a Tape station (Agilent). Libraries were sequenced using two sequencers, the Oxford Nanopore Technology (ONT) and the PacBio. Specific kits for library preparation were used whether the samples were sequenced on the PacBio or on the Oxford Nanopore Technology instruments. For the samples to be sequenced with the PacBio sequencer, the library preparation was performed using Iso-Seq Express Template Preparation for Sequel and Sequel II Systems (PacBio). For the samples sequenced with the Oxford Nanopore Technology system, the libraries were prepared using the Oxford Nanopore Technologies ligation-based kit (SQK-LSK109)-Oxford Nanopore Technologies. The library size distributions were checked using Qubit 4 (ThermoFisher) and Tapestation (Agilent).

#### Data processing, Alignment, and Differential analysis

Sequencing reads were first aligned to the human genome (GRCh38.p13) using minimap2 [29]. FeatureCounts was then used to calculate gene expression levels from transcript alignments [30]. The raw counts supplied by FeatureCounts were next provided to the R package DESeq2 [31] for differential expression analysis. Those genes with BH-FDR [32] adjusted p-values below 0.01, a log2 fold change greater than 1 or less than −1, and a base mean greater than 2 were considered differentially expressed. Genes that were considered differentially expressed using results from both of the two sequencing platforms were passed on to the subsequent GO analysis.

#### Functional annotation

The functional annotation analysis was performed using the online tool shinygo 0.75 bioinformatics.sdstate.edu/go/. [33] We focused on the GO terms that classify enrichments of differentially expressed genes according to (I) biological process, (ii) molecular function, (iii) and cellular process.

#### Enrichment analyses

We used AUTDB http://autism.mindspec.org/autdb/Welcome.do [34] to generate a list of autism-related genes to cross-reference with our lists of DEGs. We have also considered a list of FMRP targets identified in post-mortem brain tissue [35]. The enrichment analysis was performed with the following online tool http://nemates.org/MA/progs/overlap_stats.html.

## Results

### Generation of electrically active iPSC-derived cortical neurons

To evaluate neuronal development and activity, cortical neurons were first obtained from control human iPSCs (**Figure 1A**). For our protocol, we obtained cortical progenitors through neurosphere selection as described by Bell et all [23]. Once neural progenitor cells (NPCs) were obtained (**Figure 1B**), these could then be directed into cortical neurons with different neuronal differentiation media added to assess proper support for neuronal differentiation and activity *in vitro*. For this analysis, we used two different neuronal differentiation media: a supplemented serum-free neurobasal-based (NB) media and a commercial BrainPhys-based media, referred to as FB for Forebrain media (**Figure 1C**). The Neurobasal medium is widely used and generates predominantly glutamatergic neurons [23] whereas the FB media facilitates the generation of neurons from human embryonic cells and iPSC-derived precursor cells and promotes the generation of both glutamatergic and GABAergic neurons [36]. As expected, both media promoted neuronal differentiation, as we could detect the presence of the neuronal markers MAP2, VGLUT1, and GABA under both conditions (**Figure 1D-E**). Strikingly, neuronal cultures obtained with FB media presented with a more pronounced neuronal network and longer neurites as observed by MAP2 staining (**Figure 1D-E**). Neurons from NB cultures did not appear as healthy, and their processes appeared thinner. Additionally, we observed an increase in the expression of the glutamatergic neuron marker VGLUT1 in FB cultures by immunofluorescence staining (**Figure 1E**). These results indicate that FB media, a BrainPhys-based media, improved neuronal differentiation. Next, we assessed how these media types impact the activity of neurons cultivated under both conditions and to determine if neurons would be more electrically active in one of the conditions relative to the other [36].

### Multielectrode array (MEA) recording and analysis of electrical activity of iPSC-derived cortical neurons

To quantify the spontaneous neuronal activity of the neurons generated in both conditions, cells were recorded every week over a 4-week period on Multielectrode array (MEA) plates. MEA is a real-time and non-invasive technology that enables the quantification of electrophysiological function across a population of neurons simultaneously depicting information about the neuronal network [37] and its overall connectivity. As observed in our ICC tests, a neuronal network formed in both conditions albeit with more pronounced neuronal shafts in cells cultured with FB media (**Figure 2A**). Strikingly, we found an increase in spontaneous electrical activity after 3-4 weeks for both conditions across four independent experiments (**Figure 2B**). However, neurons cultivated in NB media were significantly less active relative to neurons cultivated with FB medium (**Figure 2B-D**). Neurons in FB media presented with some activity after 2-3 weeks of differentiation, whereas neurons grown in NB media had little detectable activity at the same time frame (**Figure 2B-D**). We observed a significant increase in the number of spikes (t_4weeks_=4.39, p=0.0003) and the mean firing rate (t_4weeks_=3.466, p=0.0026) in the FB media condition (**Figure 2C**). Representative raster plots showed a reduced number of spikes (black lines) in neurons cultivated with NB media (**Figure 2D**). At 4 weeks of differentiation, neurons were more active as shown in the heatmaps (**Figure 2B**), further confirmed by the increase in the number of spikes and mean firing rate (Figure 2C). Interestingly, neurons cultivated in FB medium displayed a ∼3-fold increase in their activity at 4 weeks relative to neurons maintained in NB media (**Figure 2C and D**). This demonstrates that neurons cultivated in FB media present with a higher rate of spontaneous activity, consistent with accelerated differentiation and maturation of the neurons.

**Figure 2:**
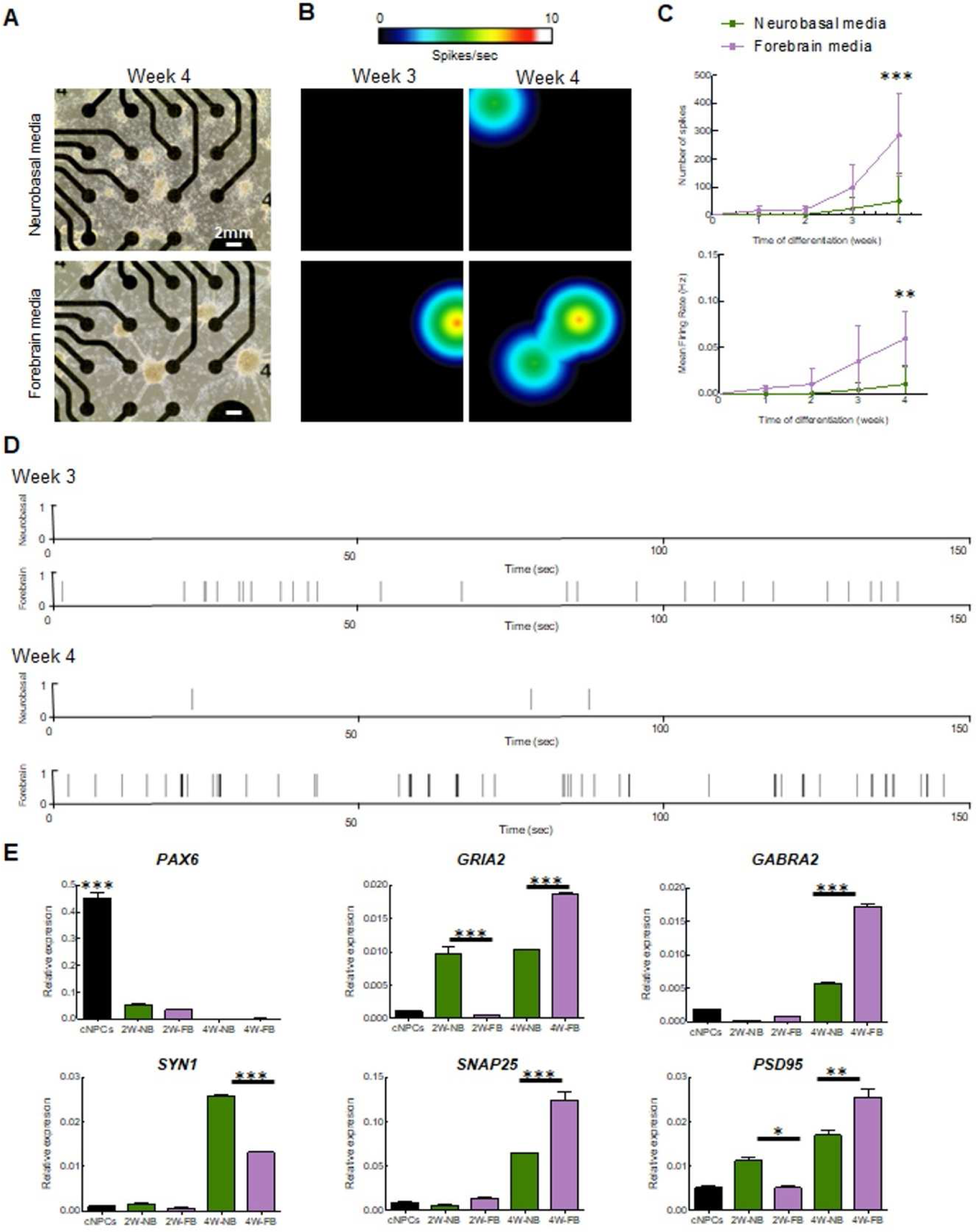
Multielectrode array (MEA) recording and analysis of electrical activity of iPSC-derived cortical neurons. (A) Light microscopy images showing cortical neurons on the MEA plate at 4 weeks of differentiation. The scale bars are 2 mm long. (B) Heat map from MEA recordings showing changes in electrical activity from week 3 to week 4 in neurons cultivated with Forebrain media. Recording analysis of cortical neurons showing (C) number of spikes, mean firing rate (C), and (D) raster plots at weeks 3 and 4 of differentiation. (E) qPCR of cortical progenitor (*PAX6*) and neuronal (*GRIA2*, *GABRA2*, *SYN1*, *SNAP25*, and *PSD95*) markers. Neurons differentiated with Neurobasal media are shown in purple, and with Forebrain media in green.

This was confirmed by examining the expression of synaptic genes in neurons cultivated with the two media (**Figure 2E**). First, we observed in both NB and FB media-treated iPSC-derived neurons a significant decrease of *PAX6* expression compared to NPCs (F=405.5; df=4; p<0.001). (**Figure 2E**) This provides support for the NPCs losing their “progenitor identity” once switched into neuronal differentiation media (either NB or FB media). For markers of neuronal differentiation, we quantified RNA postsynaptic proteins such as *SNAP25*, - Synaptosomal protein -*GABRA2-a GABA receptor subunit-*, *PSD95* – protein of the postsynaptic density-and *GRIA2 -an AMPA receptor subunit*. We observed a significant increase in the expression of both *SNAP25* and *GABRA2* mRNAs at the time of differentiation (F*_SNAP25_*=153.6, df=4, p<0.0001; F*_GABRA2_*=1651, df=4; p<0.001) and in 4W FB-differentiated neurons compared to 4W NB differentiated neurons (t*_SNAP25_* _4WNBvs4WFB_ =10.17; p<0.001); (t_GABRA2 4WNBvs4WFB_ =46.59; p<0.001) (**Figure 2E**). These findings corroborate the increased activity in 4 weeks FB media neurons suggesting that neurons in FB medium are more prone to differentiate into cortical neurons. The significant increase of *GABRA2* mRNA in 4-week-old FB neurons also reflects the ability of FB media-treated cells to commit to become GABAergic neurons. Expression levels of *GRIA2* and *PSD95*, which should be specific for glutamatergic neurons, were also significantly increased in 4 weeks FB media neurons compared to 4 weeks NB media (F*_GRIA2_*=264.4 df=4,p<0.0001; t*_GRIA2_* _2WNBvs2WFB_ =14.16; p<0.001; t*_GRIA2_* _4WNBvs4WFB_ =12.78; p<0.001; F*_PSD95_*=71,17; df=4; p<0.001 t*_PSD95_* _2WNBvs2WFB_ =4.260; p<0.05; t*_PSD95_*_4WNBvs4WFB_ =5.82; p<0.01) This shift in significant differences between the 2 weeks and 4 weeks differentiation stages could be explained by the lack of cells committed to differentiate into GABAergic neurons in the NB medium condition. We also quantified the expression levels of Synapsin1 (*SYN1)* – a presynaptic marker. Interestingly, *SYN1* expression levels were significantly higher in NB media-treated neurons compared to FB media at 4 weeks (F*_SYN1_*=2894; df=4; p<0.001; t*_SYN1_* _4WNBvs4WFB_ =43.67; p<0.001) (**Figure 2E**). This could be the result of a compensatory process of the NB medium-treated cell to balance the weak postsynaptic signaling [38].

Taken together, the MEA and qPCR results show that the FB medium is better at promoting neuronal differentiation than the NB medium. It also implies that proper differentiation of cortical neurons requires a low proportion of cells committed to becoming GABAergic neurons.

### Characterization of neuronal differentiation with *FMR1* KO iPSC

To study the impact on neuronal development when *FMR1* is absent, we generated an *FMR1* KO iPSC cell line through CRISPR/Cas9 genome editing (**Figure 3A**). CRISPR knockout cells were validated by sequencing and qPCR, and we confirmed that *FMR1* expression was abolished in the CRISPR knockout cells (**Figure 3A and E**). We successfully obtained pluripotent *FMR1* KO iPSCs as confirmed by immunohistochemistry for pluripotent markers TRA1, Nanog, SSEA, and OCT4 (**Figure 3C and D**). We assessed the expression levels of mRNA transcripts specific for the iPSC, NPCs, and neuronal stages in both the *FMR1* KO line and the isogenic control line. We quantified the iPSC-specific mRNA levels of *NANOG* and *OCT3/4* and found that their expression was significantly higher in the iPSC stage compared to NPC and Neurons (F*_NANOG_* _stage_=97.73; df=2; p<0.0001; F*_OCT3/4_* _stage_=30.67; df=2; p<0.0001) (**Figure 3E**). These findings confirm the results collected from the immunostaining performed in iPSCs (**Figure 3C-D**). They also demonstrate that knocking out the *FMR1* gene does not affect the pluripotency of iPSCs. Moreover, the absence of iPSC markers at the NPC and neuronal stages shows that *FMR1* KO cells can be induced toward neuronal subtypes. As mentioned previously, the *FMR1* KO was generated by deleting part of the exon 5 of the *FMR1* gene sequence. We quantified *FMR1* mRNA expression level at the iPSC, NPC, and neuron stages and found decreased *FMR1* levels in *FMR1* KO iPSC that became significant in *FMR1* KO NPCs and neurons (F*_FMR1_* _stage_=79.45; df=2; p<0.0001; t_NPC_=22.95; p<0.0001; t_Neurons_=17.97; p<0.0001) (**Figure 3E**). This demonstrates that the deletion in exon 5 reduced the transcription of the *FMR1* gene. Next, with neural progenitors and cortical neurons generated from the *FMR1* KO isogenic line, we tested for neuronal differentiation. Intriguingly, morphological alterations were observed in neuronal cells derived from the *FMR1* KO cells. Although the *FMR1* KO cells were able to produce progenitors, as confirmed by the presence of the progenitor cell marker Nestin (**Figure 4A**, left panels), we observed a drastic reduction of the neuronal population relative to the isogenic control. This was confirmed by our findings that few Tuj1 and MAP2 positive *FMR1* KO neurons were detected after 3 or 4 weeks of differentiation (**Figure 4A** right panels) compared to the isogenic control neurons. Neuronal differentiation was also altered when *FMR1* was absent, as *FMR1* KO neurons displayed an altered morphology with shorter processes compared to control neurons. Taken together, disruption of the neuronal network was evident in *FMR1* KO lines at 4 weeks of differentiation with a significant reduction of the neuronal population (**Figure 4A**, left panels).

**Figure 3:**
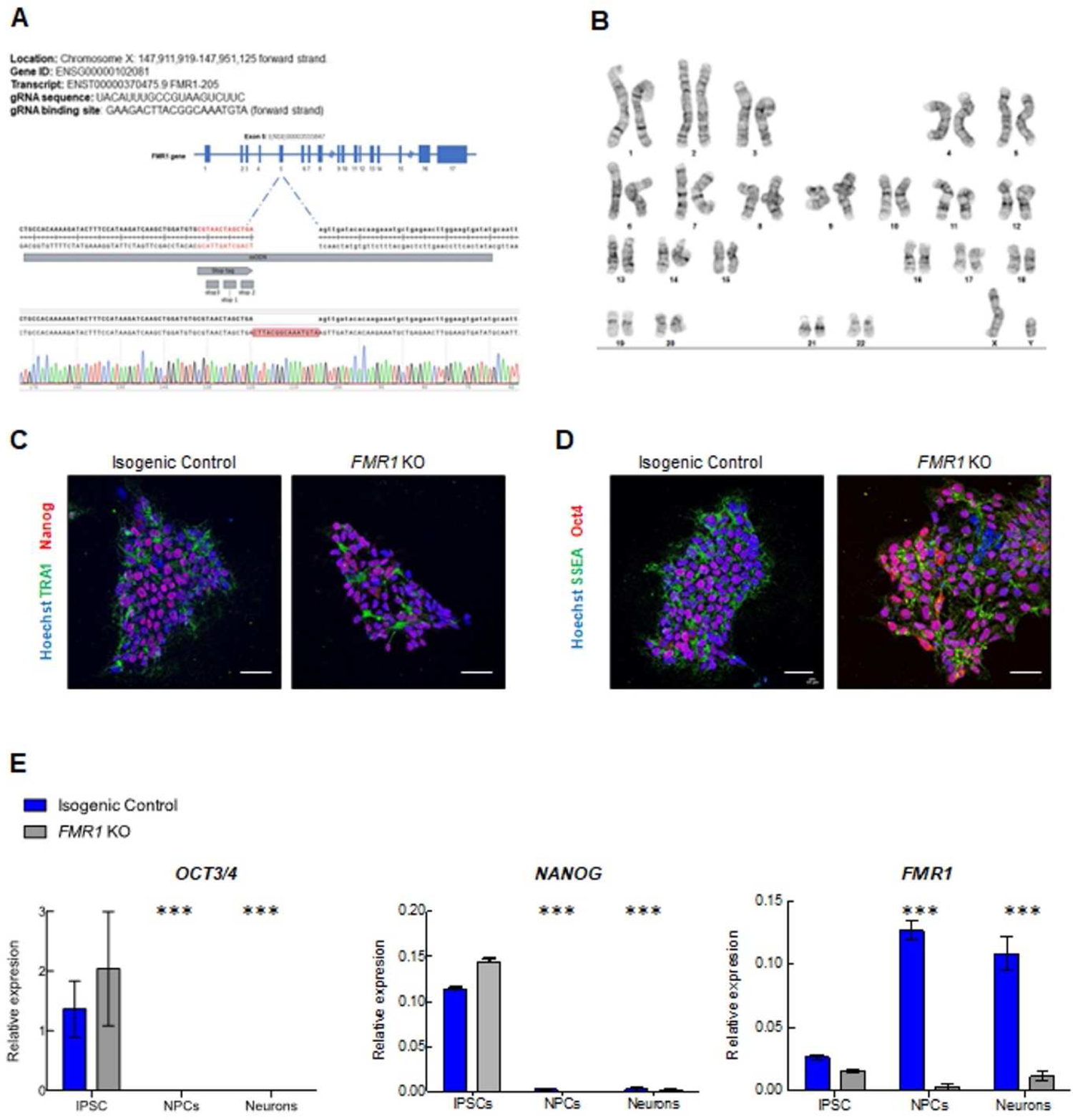
Generation of *FMR1* KO iPSC and neural progenitors. (A) Validation of CRISPR/Cas9 editing of the *FMR1* gene: *FMR1* KO nucleotide sequencing results showing insertion of stop tag in exon 5 of the *FMR1* gene. (B) Karyotyping of *FMR1* KO. G-banding chromosome analysis showed Normal 46 XY. (C-D) immunofluorescence images of isogenic control and *FMR1* KO iPSCs showing expression of pluripotent markers (TRA1, Nanog, SSEA and OCT4). Nuclei were counterstained with Hoechst. The scale bars are 50 µm. (E) qPCR expression of the pluripotent genes *OCT3/4* and *NANOG*, and *FMR1* in iPSC, NPCs, and 3 weeks cortical neurons.

**Figure 4:**
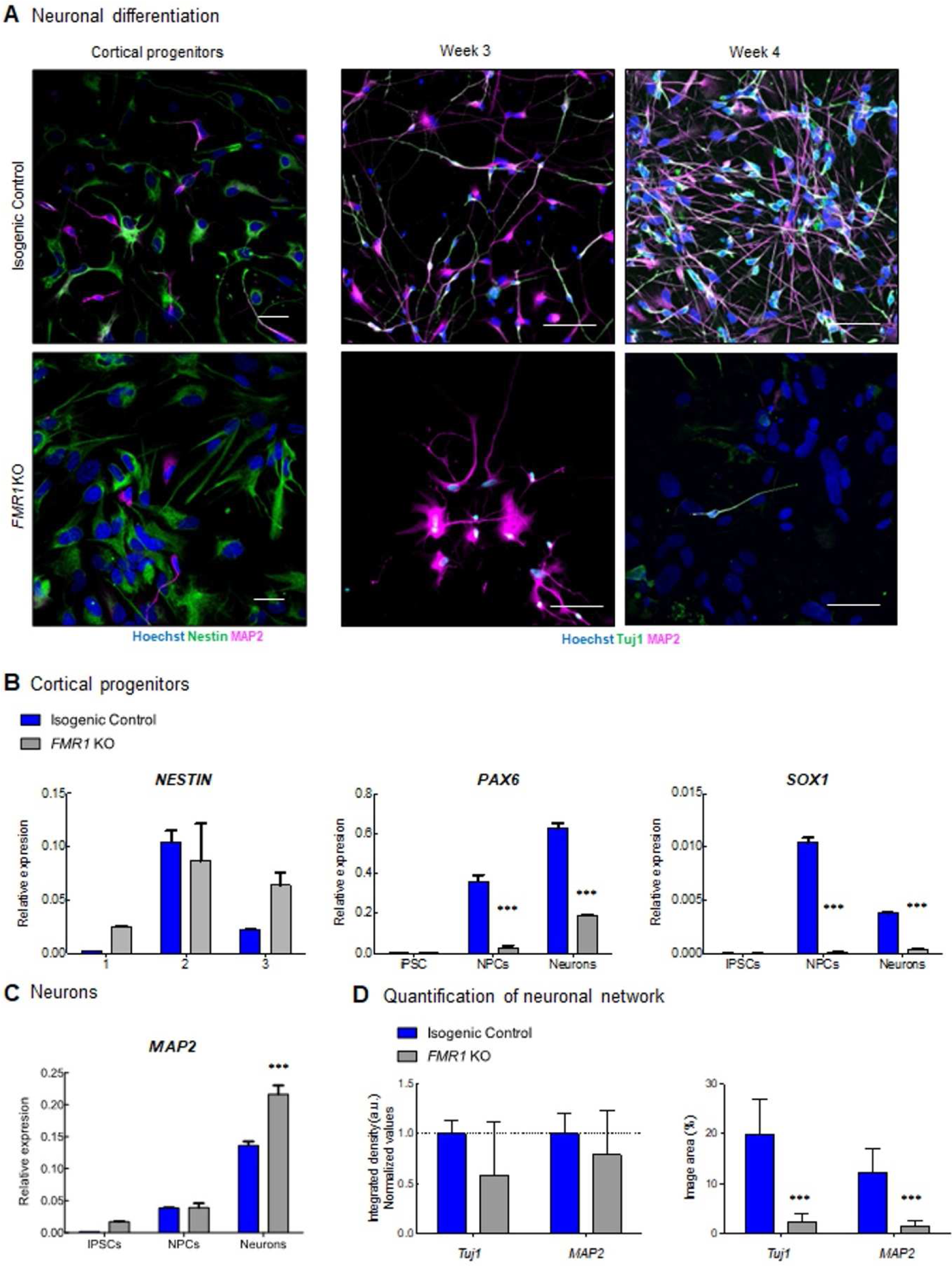
Altered neuronal development in *FMR1* KO. (A) Immunofluorescence images of cortical progenitors and cortical neurons at 3 and 4 weeks of differentiation showing expression of progenitor markers Nestin (green), and neuronal markers (Tuj1 and MAP2). Nuclei were counterstained with Hoechst. The scale bars are 50µm. qPCR expression of (B) neuronal progenitors (*SOX1*, *NESTIN* and *PAX6*) and (C) neuronal (*MAP2*) genes. (D) Quantification of neuronal network showing integrated density (a.u.) for Tuj1 and MAP2 at 4 weeks of differentiation.

Next, we quantified the expression of mRNA specific to NPC and neuronal stages to determine if the knockout of *the FMR1 gene a*ffected the identity of the progenitor population (**Figure 4B and C**). First, significant changes in *Nestin* expression were observed across stages (F*_Nestin_* _stage_=12.88; df=2; p=0.0013) with higher expression in control NPCs compared to iPSCs (t=4.427; p=0.001) and neurons (t=3.162; p=0.009), as well as higher expression in *FMR1* KO NPCs compared to *FMR1* KO iPSCs (t=2.677; p=0.0215) but not neurons (t=0.9828; p=0.3468) (**Figure 4B**). PAX6 expression levels increase from the iPSC to the neuron stages (F*_PAX6_* _stage_=197.2; df=2; p<0.0001) with significantly lower levels observed in *FMR1* KO NPCs and Neurons (F_PAX6 genotype_=243.7; df=1; p<0.0001; t_NPCs_=12.10; p<0.001; t_Neurons_ =14.29, p<0.001)) Finally, we observed change in *SOX1* expression across all stages (F SOX1 Stage=32.05; df=1; p<0.0001) with significantly higher levels in control NPCs (t=35.44; p<0.001) and neurons (t=11.69;p<0.001) compared to *FMR1* KO cells. To assess the ability of the iPSCs to differentiate into neurons, we quantified *MAP2* expression as a neuronal marker in iPSCs, NPCs, and neurons. Using two-way ANOVA, we observed a significant increase in *MAP2* expression from the IPSC to the neuronal stage (F*_MAP2_* _stage_=331.8; df=2; p<0.0001) (**Figure 4C**). At the neuronal stage, a significantly higher *MAP2* mRNA expression level is detected in *FMR1*-KO neurons compared to isogenic control neurons (t*_MAP2_* _Neurons_ =8.207; p<0.001). This increase corroborates the intensity levels captured by immunostaining (**Figure 4A**). These findings demonstrate that neurons can be derived from the *FMR1*-KO cell line. However, knocking out the *FMR1* gene affects (i) the cellular identity of the progenitor and (ii) the capability of the cells to form a neuronal network.

### Neuronal spontaneous electrical activity impairment in the *FMR1* KO

Given our earlier findings in which the neuronal network was disrupted when *FMR1* was knocked out, we next assessed the activity of neurons that could form from *FMR1* KO cells by MEA, and as expected it was significantly reduced in comparison to isogenic control cells (**Figure 5**). As outlined above, we plated neuronal progenitor cells in MEA plates (**Figure 5A**) and measured the spontaneous activity at weekly intervals over a 4-week period. We observed an increase in neuronal activity around week 3 in the isogenic control neurons that was even more pronounced at week 4 (**Figure 5B-E**). As predicted, we observed impairment in the spontaneous activity in *FMR1* KO neurons, as depicted in the heatmaps, with few spots of activity detected (**Figure 5B**), and consequently a lower number of spikes (**Figure 5C and D**) and reduced mean firing rate (**Figure 5E**). Electrical activity, after treatment with voltage-gated sodium channel blocker Tetrodotoxin (TTX) (**Figure 5F**) was abolished (F=52.34,df=1,p<0.0001), leading to inhibition of neuronal transmission, indicating that neuronal activity results from sodium currents [39]. This analysis showed that the absence of FMRP disrupts neuronal maturation and electrical activity in our model.

**Figure 5:**
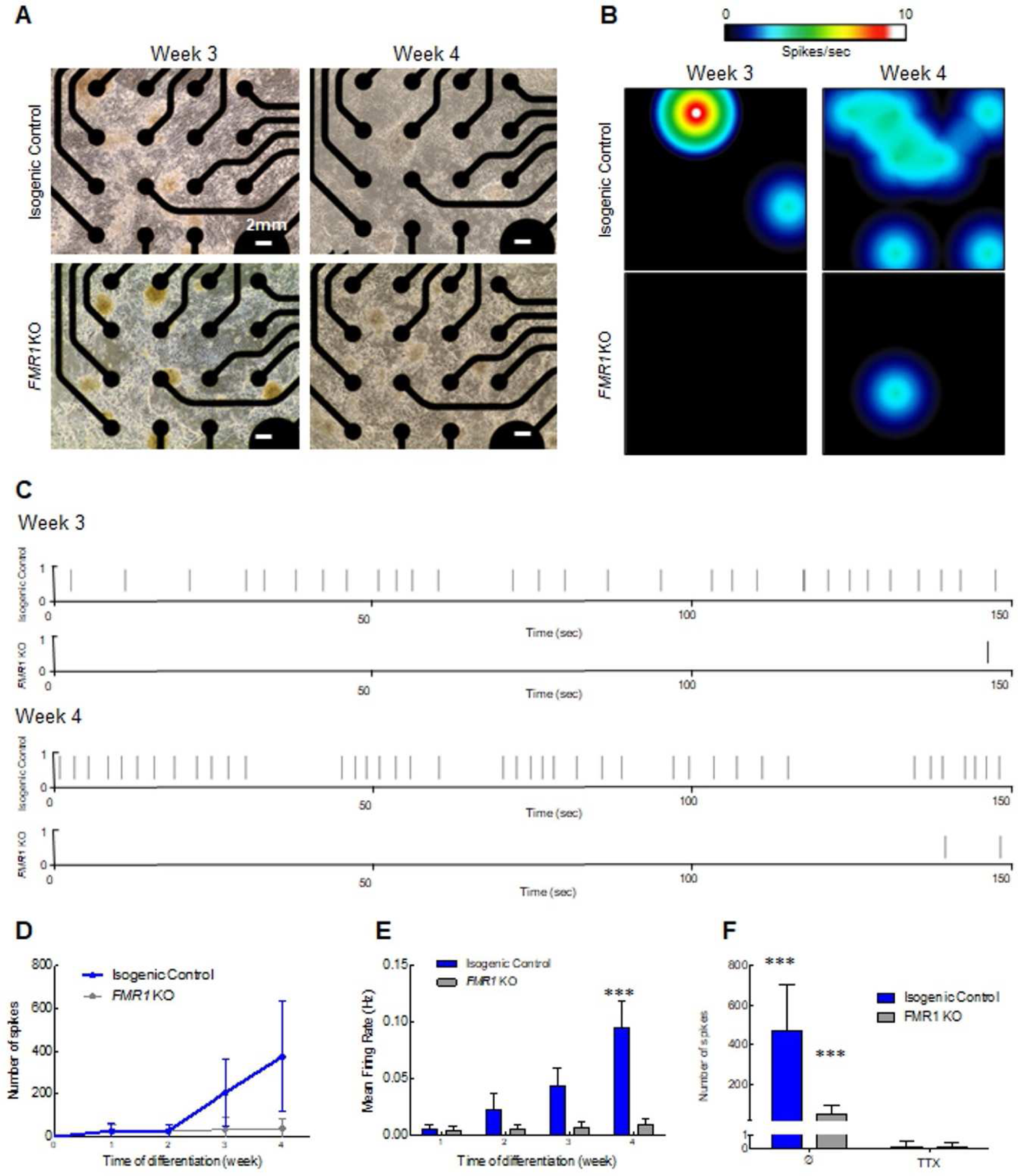
Neuronal spontaneous electrical activity impairment in the *FMR1* KO. (A) Light microscopy images showing cortical neurons on the MEA plate at 3 and 4 weeks of differentiation. The scale bars are 2 mm long. (B) Heat map from MEA recordings showing changes in electrical activity from week 3 to week 4 in neurons. Recording analysis of cortical neurons showing (C) raster plots from weeks 3 and 4, (D) number of spikes, and (E) mean firing rate. (F) Number of spikes after 30 min of 1 µm TTX treatment.

### Whole transcriptome profiling of *FMR1* KO iPSC-derived NPCs and neurons

To better investigate and understand the molecular changes induced by suppressing the *FMR1* gene expression in iPSC-derived cells, we analyzed the whole genome expression profile of iPSC-derived *FMR1*-KO NPCs and iPSC-derived neurons in comparison with their respective isogenic control cells using an RNA sequencing approach. We also analyzed the common deregulations between *FMR1* KO NPCs and *FMR1* KO neurons (**supplementary information** and **supplementary figure 2**). At the NPC stage, we found 1,316 genes differentially expressed between *FMR1* KO and isogenic control cell lines, (738 downregulated and 578 upregulated in *FMR1* KO; **Figure 6A**). 1,307 genes were also found to be differentially expressed between *FMR1* KO iPSC-derived neurons and controls (789 genes were found downregulated and 518 upregulated; **Figure 6B**). The analysis featuring the sets of genes that are commonly deregulated in *FMR1* KO NPCs and neurons is presented in **supplementary figure 2** and **supplementary information**.

**Figure 6:**
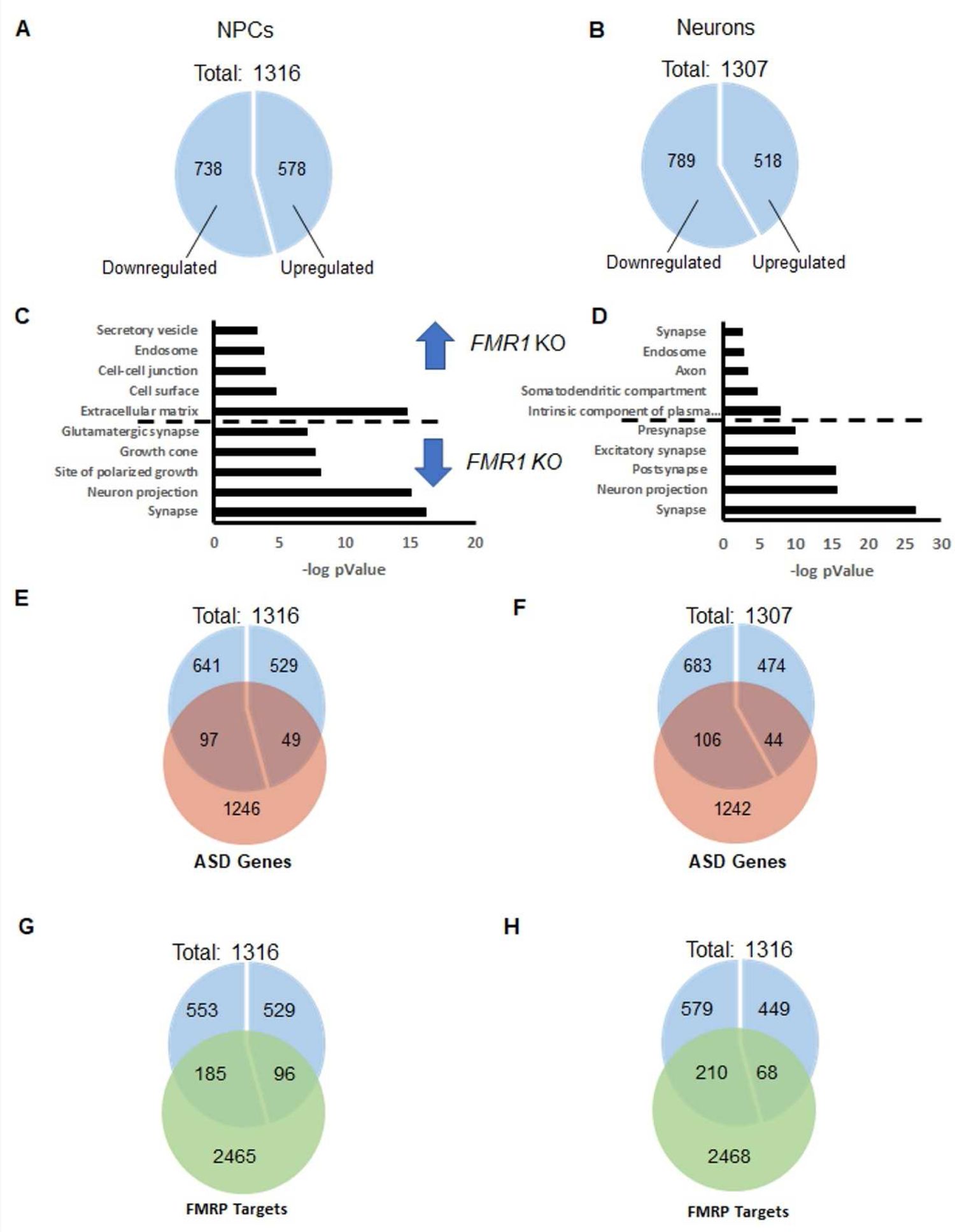
Whole transcriptome profile of *FMR1* KO iPSC-derived NPCs and neurons. Fractions of genes upregulated and downregulated in A) the *FMR1* KO NPCs and (B) the *FMR1* KO iPSC-derived neurons. (C) and (D) Functional annotation analysis presenting the GO terms with significant enrichment of the differentially expressed genes in iPSC-derived NPCs and IPSC-derived neurons respectively. GO enrichment for upregulated and downregulated genes separated by a dotted line. (E) and (F) Overlaps between autism risk genes and the differentially expressed genes in iPSC-derived *FMR1* KO NPCs, and *FMR1* KO neurons, respectively. (G) and (H) Overlaps between the list of FMRP targets and differentially expressed genes in the *FMR1* KO iPSC-derived NPCs and the *FMR1* KO iPSC-derived, respectively.

The functional annotation analysis based on GO Cellular Component Terms shows that many genes upregulated in *FMR1* KO NPCs are related to the extracellular matrix and the cell-cell junction. Interestingly, genes downregulated in *FMR1* KO NPCs are associated with the synapse, neuronal projection, and growth cone processes (**Figure 6C**). This information shows that synaptic genes are already expressed at the NPC stage and implies that *FMR1* KO NPCs may be less prone to differentiate into neurons. Genes upregulated in *FMR1* KO neurons are involved in the intrinsic component of the plasma membrane, in the somatodendritic compartment, in the axon, and in the endosome.

Interestingly, we found, that significantly downregulated genes in *FMR1* KO neurons are related to the synapse, and more precisely to the glutamatergic excitatory synapse and neuronal projections. The significant enrichment, in *FMR1* KO NPCs and *FMR1* KO neurons, of downregulated genes involved in the synapse, combined with an enrichment of upregulated genes associated with the plasma membrane in *FMR1* KO neurons, implies that those differentiating cells are being maintained in a progenitor-like state with impaired differentiation into neurons.

As fragile X syndrome is considered a syndromic autism, we next investigated whether autism-related genes and FMRP targets are overrepresented in our lists of differentially expressed genes. We have separately considered genes that were down or upregulated in *FMR1* KO iPSC-derived cells. We used the list of Autism Related Genes established by the database AutDB [34] and the list of FMRP targets as our reference. We also compared our lists to the list of FMRP targets [35].

From these lists, we observed a significant enrichment of autism-related genes (97/738; p < 4.004e-07; **Figure 6E**) and FMRP targets (185/738, 6.885e-12) (**Figure 6G**) in the list of genes downregulated in *FMR1* KO NPCs, but not in the list of genes that were upregulated in *FMR1* KO NPCs. Similar enrichment in autism-related genes (106/789; p < 3.680e-08 **Figure 6F**) and FMRP targets (210/789; p < 3.056e-16 **Figure 6H**) was identified from the list of genes downregulated in *FMR1* KO neurons. Taken together, this data confirms that there are shared, commonly deregulated pathways and differentiation processes in idiopathic autism and Fragile X syndrome. These common deregulations include and are likely driven by a reduced expression in specific synaptic genes.

Our analyses (**Figure 6** and **Supplementary** Figure 2) also indicate that a substantial proportion of differentially expressed genes are deregulated at the NPC and neuronal stages and that a considerable proportion of downregulated genes in *FMR1* KO cells includes autism-related genes and FMRP targets. This also confirms that changes of expression in genes initially described as involved in the synapse might be deleterious to the cell development and function at early stage of the neurogenesis.

### Differential expression of synaptic markers in the *FMR1* KO

In assessing the activity of *FMR1* KO iPSC-derived cells, we observed a significant decrease in the mean firing rate of these cells when differentiated for 4 weeks (**Figure 5**). Interestingly, the whole transcriptome expression profile revealed a significant enrichment of genes associated with the synapse and whose expression levels were decreased in *FMR1* KO iPSC-derived cells (**Figure 6**). Thus, we hypothesized that decreased expression of synaptic molecules affects the activity of *FMR1* KO differentiating cells. We quantified *SYN1*, *SYP*, and *SLC17A7* mRNA which are coding presynaptic proteins. At the NPC stage, no significant change of expression was observed between *FMR1* KO cell lines and isogenic control cell lines (t*_SYN1_* _NPCs_=1.401; p>0.05; t*_SYP_* _NPCs_=0.3670; p>0.05; t*_SLC17A7_* _NPCs_=0.06409; p>0.05) (**Figure 7A**). At the neuronal stage, we found a significant increase in the expression of the three transcripts in the *FMR1* KO line (t*_SYN1_* _Neurons_=3.011; p<0.05; t*_SYP_* _Neurons_=7.527; p<0.001; t*_SLC17A7_* _Neurons_=7.913; p<0.001) (**Figure 7A**). We also assessed the expression of the following postsynaptic mRNA *GRIA1* and *GRIA2* (coding for AMPA receptor subunits); *NTRK2* (coding for BDNF receptor), *GRIN2B* (coding for an NMDA receptor subunit), *GABRA2* (coding for a GABA receptor subunit) and *SNAP25* (Synaptosomal protein) No significant changes were observed in the *GRIA1* (t*_GRIA1 NPC_*_s_=0.2744; p>0.05), *GRIA2 (t _GRIA2 NPC_*_s_ *=0.009397; p>0.05)*, *SNAP25* (t*_SNAP25 NPC_*_s_ =0.7854; p>0.05) *GRIN2B* ((t*_GRIN2B NPC_*_s_ =0.9188; p>0.05) and *GABRA2 (t_GABRA2 NPC_*_s_ *=0.1113; p>0.05)* expressions at the NPC stage (**Figure 7A**) between control and *FMR1* KO line. Nevertheless, we validated that for these five transcripts, there was a significant decrease in expression for each of these five genes in the *FMR1* KO neurons compared to isogenic control (t*_GRIA1 Neurons_*=8.109; p<0.001), *GRIA2 (t_GRIA2 Neurons_=11.14; p<0.001)*, *GRIN2B* (t=5.718; p<0.001) *SNAP25* (t*_SNAP25 Neuron_*_s_ =4.958; p<0.01) and *GABRA2 (t_GABRA2 Neuron_*_s_ *=17.69; p<0.001*. We also demonstrate that expression of the *NTRK2* gene was significantly decreased in the *FMR1* KO cells at both stages relative to isogenic controls (t*_NTRK2_* _NPCs_=9.770, df=8, p<0.0001; t*_NTRK2_* _Neurons_=38.22, df=8, p<0.0001; (**Figure 7A**). Strikingly, low levels of SLC17A7/VGLUT1, a glutamate transporter that carries glutamate to synaptic vesicles, were observed by immunofluorescence at 4 weeks of differentiation in isogenic control cell lines, whereas higher VGLUT1 signal was detected in *FMR1* KO neurons confirming our qPCR results (**Figure 7B**).

**Figure 7:**
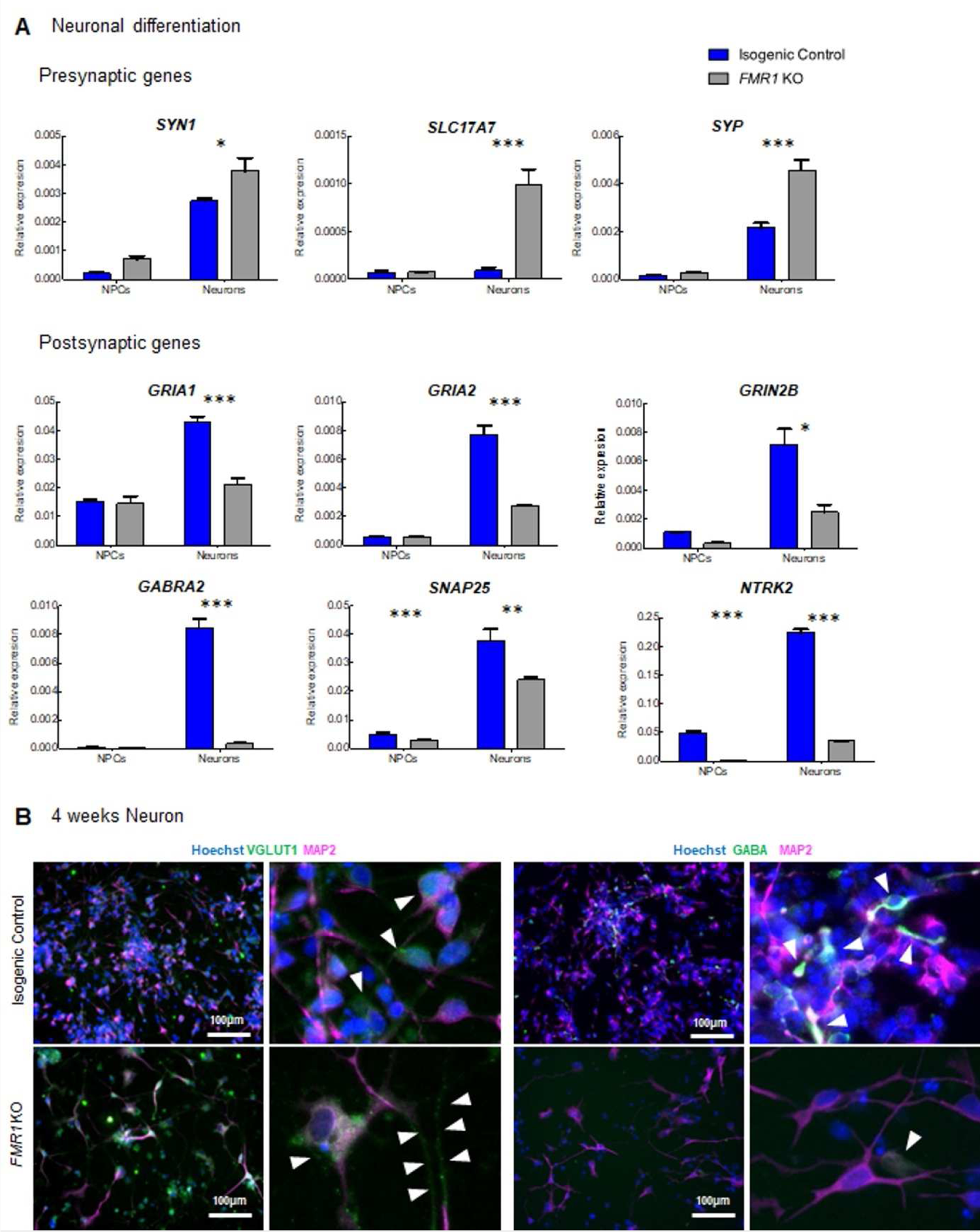
Differential expression of synaptic markers in the *FMR1* KO. (A) qPCR expression of the presynaptic genes *SYN1*, *SLC17A7*, *SYP*, and the postsynaptic genes *GRIA1, GRIA2, NTRK2, GRIN2B, GABRA2* and *SNAP2*5 in neuronal progenitors and neurons from isogenic control and *FMR1* KO cell lines. (B) Immunofluorescence images of 4 weeks cortical neurons. Images show synaptic markers VGLUT1 and GABA (green) and MAP2 (magenta). Nuclei were counterstained with Hoechst. The scale bars are 100 µm.

We were also able to detect neuronal processes enriched in VGLUT1 positive particles in *FMR1* KO neurons as shown in **Figure 4B**. However, we did not observe the same alterations in the GABAergic-associated proteins as expression levels for the GABRA2 receptor were downregulated in *FMR1* KO neurons and, similarly, GABA+ neurons were not detected by immunofluorescence (**Figure 4B**). Thus, the decreased expression of postsynaptic molecules would appear to contribute towards the decreased mean firing rate observed with MEA. On the other hand, the increase of presynaptic molecules such as VGLUT1 could represent a process to compensate for the low levels of postsynaptic molecules.

### Fragile X syndrome patient cell line displays impaired neuronal development and activity

Lastly, since we established (1) a protocol to obtain electrically active iPSC-derived cortical neurons in 4 weeks of differentiation in vitro, (2) created a pipeline for neuronal development analysis from quality control to neuronal activity assessment by MEA, (3) demonstrated that *FMR1* KO neurons display altered neuronal development and impaired neuronal activity *in vitro* (**Figure 9**), Thus, next we sought to evaluate neuronal differentiation and spontaneous activity in the, a Fragile X Syndrome Patient cell line (referred to as FX11-7) relative to a control line. Consistent with our findings in an *FMR1* KO cell line, progenitors from the FX11-7 cell line exhibited altered morphology with a more elongated and slender profile when compared to a control cell line (**Figure 8A** upper panels). This altered morphology in the progenitor population showed an impairment in the early stages of neuronal development. To study neuronal differentiation in FXS cells, we applied the culture protocol described in **Figure 1 and 2** using FB media. Although neurons could form, as seen by the presence of Tuj1 and MAP2 positive cells, patient cells displayed impairment in neuronal development as a reduced number of neurons were found in these cultures (**Figure 8A**, lower panels).

**Figure 8:**
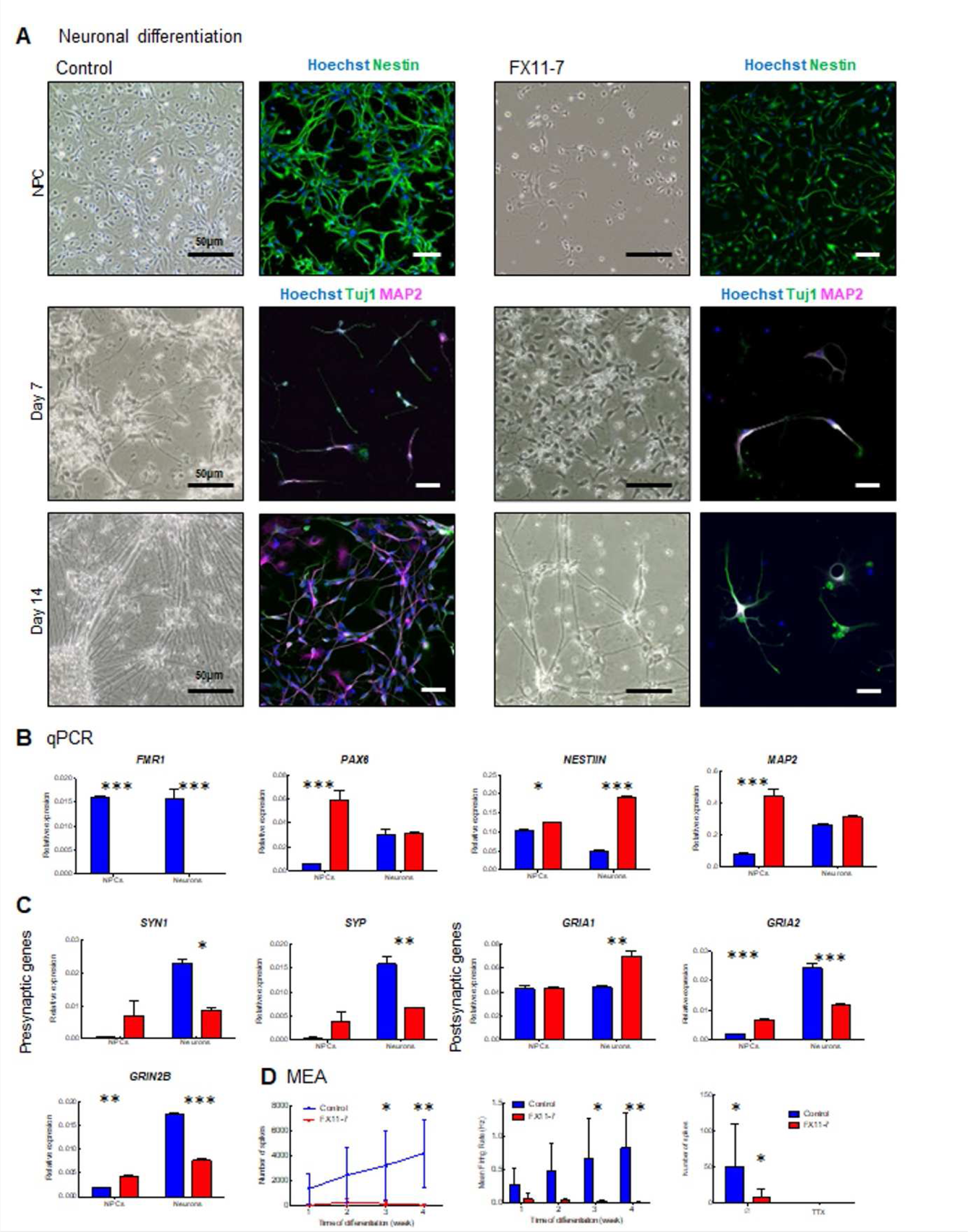
Fragile X syndrome patient cell line displays impaired neuronal development and activity. (A) Light microscopy and immunofluorescence images of cortical progenitors at day 7 and day 14 cortical neurons. Images show the neuronal progenitor marker Nestin (green), and the neuronal markers Tuj1 (green) and MAP2 (magenta). Nuclei were counterstained with Hoechst. The scale bars are 50 µm. (B) qPCR expression of *FMR1*, progenitor (*PAX6* and *NESTIN*) and cortical neuron (*MAP2)* genes. (C) qPCR expression of presynaptic (*SYN1*, *SYP*) and postsynaptic (*GRIA1*, *GRIA2*, and *GRIN2B*) genes in neuronal progenitors and cortical neurons from control and FX11-7 cell line neurons. (D) MEA analysis showing number of spikes and mean firing rate, and number of spikes after 30 min of 1 µm Na+ channel blocker TTX treatment.

**Figure 9:**
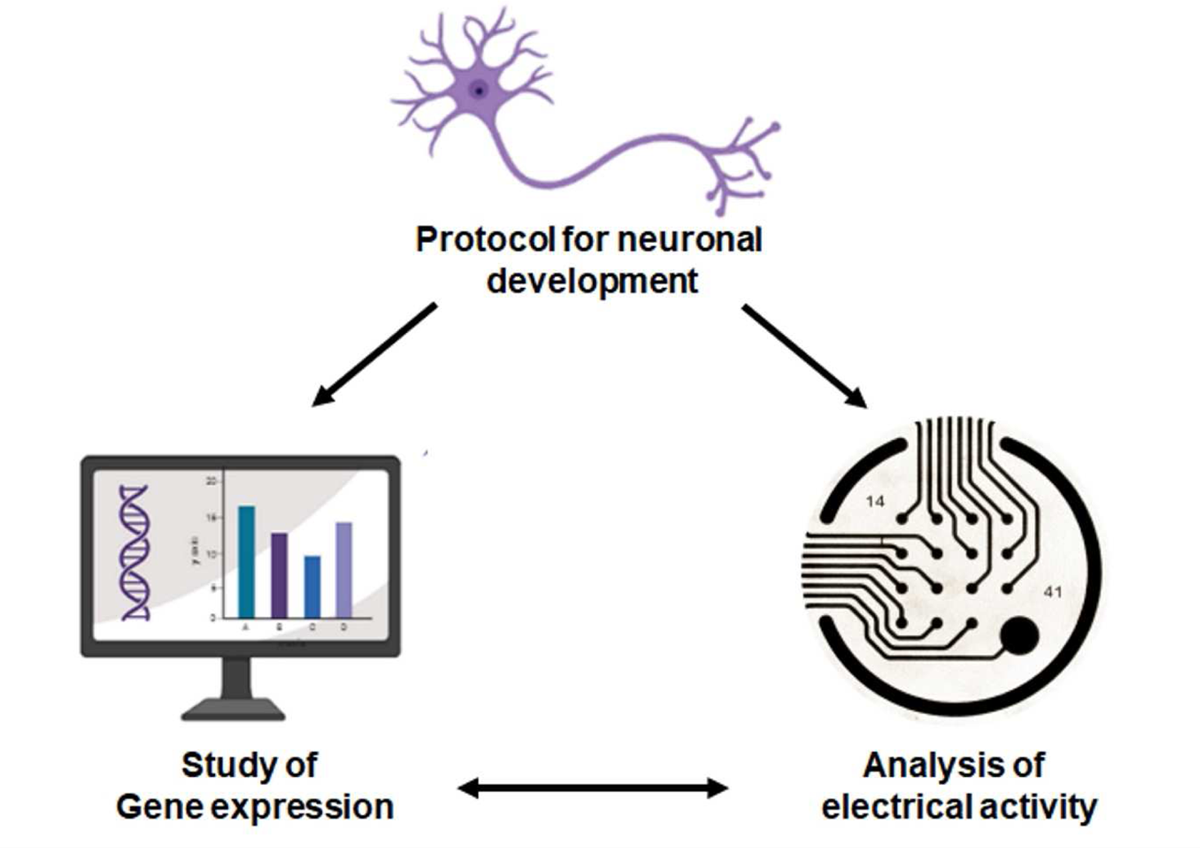
Neurodevelopmental study pipeline. Workflow for the generation of electrically active iPSC-derived cortical neurons, assessment of neuronal activity by MEA, and analysis of the expression of genes involved in synaptic transmission.

Building on these findings, we investigated the expression profiles of synaptic genes and the activity of iPSC-derived cells from patients diagnosed with Fragile X compared to a control line. First, assessing the *FMR1* gene expression in both lines, no expression was observed in neurons derived from the patient line. That result confirms that the CGG expansion in the 5’UTR sequence of the *FMR1* gene in the patient cells completely silences its expression (**Figure 8B**). Markers of neuronal progenitors (*Nestin* and *PAX6*) and neurons (*MAP2*) were also quantified. For these three transcripts, using a two-way ANOVA, we found significant interactions between genotype and differentiation stage for their expression levels (F*_NESTIN_* _GenotypeXStage_=213.7 df=1 p<0.0001; F*_PAX6_* _GenotypeXStage_=48.03 df=1 p=0.0004; F*_MAP2_* _GenotypeXStage_=64.25 df=1 p<0.0002). Although expected, in control condition, *Nestin* expression is decreased at the neuron stage compared to NPCs. Moreover, we observed a significant increase in *Nestin* levels in FXS NPCs (t=3.338 p<0.05) and Neurons (t=24.01 p<0.001) compared to control. A significant increase in *PAX6* expression was observed in FXS NPCs (t=10.05 p<0.001) but not in neurons (t=0.2534 p>0.05) compared to control cells. Finally, we observe a significant increase of *MAP2* expression in FXS NPCs (t=13.21, p<0.001) compared to controls but not in neurons (t=1.872; p>0.05). Combined, those results suggest that FXS iPSC-derived neurons are maintained in a progenitor-like state.

Similar to earlier work in the *FMR1* KO, we next quantified mRNA coding presynaptic proteins (Synaptophysin-*SYP*, and Synapsin I-*SYNI)*, and mRNA coding post-synaptic proteins (*GRIA1*; *GRIA2* and *GRIN2B*) (**Figure 8C**). Interestingly, except for *GRIA1* expression levels, similar patterns were found between the four other transcripts. We observed significant interactions between genotypes and stage for *SYN1* (F*_SYN1_* _GenotypeXStage_=14.13 df=1 p=0.0071); *SYP* (F*_SYP_* _GenotypeXStage_=23.51 df=1 p=0.0019); *GRIA2* (F*_GRIA2_* _GenotypeXStage_=27.66 df=1 p<0.0001) and *GRIN2B* (F*_GRIN2B_* _GenotypeXStage_=492.7 df=1 p<0.0001) expressions with, compared to the respective controls; increases in FXS NPCs (t*_SYN1_*=1.713 p>0.05; t*_SYP_*=1.893 p>0.05; t*_GRIA2_*=4.826 p<0.001; t*_GRIN2B_*=6.129 p<0.01) and significant decreases in FXS neurons compared (t*_SYN1_*=3.511 p<0.05; t*_SYP_*=4.812 p<0.01; t*_GRIA2_*=12.48 p<0.001; t*_GRIN2B_*=25.26 p<0.001). Comparing specific gene expression levels in FXS patients’ lines and control; we could partially duplicate findings from the *FMR1* KO line. Interestingly, the increased expression of synaptic molecules at the NPCs stage may be explained by a lack of transcriptional regulation of FMRP targets in FXS cells. At the neuronal stage, the decreased expressions of *GRIA2* and *GRIN2B* also observed in the *FMR1* KO line, may reflect, an impaired neuronal differentiation and/or an increased proportion of differentiating non-neuronal cells in the FXS cells.

Next, we checked the neuronal activity of neurons from the patient line, and we found, similar to the *FMR1* KO cell line that FX11-7 neurons display a low level of activity throughout the period of 4 weeks of neuronal differentiation when compared to controls. A significant decrease in the number of spikes with patient cells (F_genotype_=15.65; df=1,p=0.0011) at 3 weeks (t=2.270 p=0.0324 and 4 weeks of differentiation (t=3.061, p=0.0075), with no progressive increase in activity over time (F_Stage_=0.8002; df=3; p= 0.5118; **Figure 8D**) was also observed. That observation also translated into a significant decrease in the mean firing rate in neurons from our patient line compared to the control (F=15.64, df=1,p=0.0011) with a significant reduction at 3 weeks (t=2.403,p=0.0288) and 4 weeks of differentiation (t=3.053, p=0.0076). Finally, cells were treated with TTX which abolishes, the activity in both control and patient lines (F=7.015, df=1, p=0.0119). This shows that altered progenitor population and impaired differentiation directly contribute to neuronal activity deficits in our model.

While common changes were observed in terms of activities and gene expression between *the FMR1* KO line and the FXS patient cell line, we also assessed the activity of neurons from our patient line using a calcium imaging approach. As we found a significant decrease in the expression of *GRIN2B* expression in the FXS patient neurons (also observed in the *FMR1* KO neurons), we tested the NMDA response on the FXS patient cells compared to controls. We found a significant decrease in the NMDA response, at the NPC stage (t_NPCs_=45.87; df=699; p<0.0001) and at the neuronal stage (t_Neurons_=8.060; df=711; p<0.0001), compared to our control. These findings imply that the absence of FMRP protein, affects the synthesis of NMDA receptor subunits, contributing to calcium influx in the cells that is necessary for proper neuronal differentiation in the early stages, although synapses are not formed yet.

Taken together, our study highlights the importance of optimizing differentiation protocols of iPSC-cortical neurons for investigating neuronal dysregulation in the context of neurodevelopmental disorders. With this protocol established, we could next define specific phenotypes that were arising- and in particular decreased activity and expression of synaptic genes in both the *FMR1* KO and Fragile X patient lines. These findings suggest that an absence of FMRP protein affects the activity-dependent development of the cells at early stages of differentiation, preceding synapse formation.

## Discussion

In this study, we developed an *FMR1* KO iPSC-derived model to better understand how suppressing the expression of the *FMR1* gene contributes to neurodevelopmental alterations that are observed in Fragile X syndrome. We applied the most efficient protocol for cortical differentiation to investigate differences in activity and gene expression in *FMR1* KO cells derived from CRISPR–edited iPSC and their endogenous controls. While *FMR1* KO neurons exhibit a decreased activity compared to the controls, we also show that *FMR1* KO NPCs and neurons share 40% of differentially expressed genes. In the genes that are down-regulated, significant enrichment was observed (i) in genes involved in autism-related genes, (ii) in FMRP targets, and (iii) in synaptic function. These findings imply that, although the synapses are not formed yet, synaptic gene products (including scaffolding protein and receptor subunits), are required for proper neuronal differentiation. We also found expression of presynaptic molecules to be increased by RNA sequencing, qPCR, and immunostaining. Taken together, observations in both iPSC-derived FXS patient cells and *FMR1* KO line are suggestive of neuronal differentiation being impaired at the initial stages of neurodevelopment.

### FMRP, a multifunction protein

FRMP has initially been described as an RNA-binding protein that acts as a translational repressor [40]. More recent studies have shown that FMRP also regulates (i) the expression of chromatin modifiers and (ii) the activity of transcription factors and channel receptors through protein-protein interaction [3, 4]. Interestingly, in our RNA sequencing experiment, we found approximately 1300 differentially expressed genes in both the NPCs and neurons, with 55% of them downregulated and 45% upregulated. We also observed that of these differentially expressed genes, the FMRP targets are not restricted to genes that are upregulated in *FMR1* KO cells. This confirms that FMRP does not only act as a translational repressor. More precisely, in *FMR1* KO cells we observed an upregulation of transcription factors such as *SIX3* and *FOXG1 and* three presynaptic mRNAs *SLC17A7 SYP* and *SYN1*, having been described as FMRP targets [35]. On the other side, we have found the expression of *GRIA2* and *GRIN2B* to be decreased, with each coding AMPA and NMDA receptor subunits respectively. The proportion of upregulated and downregulated genes in *FMR1* KO cells; as well as their functions reflects the different mechanisms through which FMRP can modulate gene and protein expression.

### Delayed transition from progenitor-like state towards a neuron and impaired differentiation

iPSCs from the *FMR1* KO and the isogenic control lines were induced into NPCs that then went through cortical differentiation. In the *FMR1* KO NPCs population, we observed a significant enrichment of upregulated genes involved with extracellular matrix combined with the downregulation of synaptic genes. Inversely, synaptic genes are downregulated in *FMR1* KO neurons. These results suggest that *FMR1* KO NPCs are less prone to differentiate into cortical neurons and/or *FMR1 K*O NPCs are being maintained in a progenitor-like state. As shown previously by Raj et al., 2021 [41], we also observe impaired cell fate specification that favors proliferative over neurogenic cell fates during development. The functional annotation performed on genes commonly deregulated at the NPC and neuronal stages has revealed the upregulation of genes involved in forebrain development. As such, we found a significant increase of *SIX3* and *FOXG1* gene expression in *FMR1* KO NPCs and neurons when compared to control neurons. Both genes are transcription factors involved in the specification of the telencephalon brain region. Interestingly, according to data collected by the brain span consortium (http://www.brainspan.org/), the peak of expression of *SIX3* is located around 8 weeks post conception but its expression remains in the amygdala and the striatum whereas the *FOXG1* gene, which is more expressed throughout different brain regions, remains predominant in cortical areas but its expression is decreased with age. The increase in expression of both transcription factors, combined with the decreased expression of synaptic genes in *FMR1* KO cells, suggests that in the absence of FMRP, neural progenitors are less committed to differentiate into cortical neurons and the neural cell fate remains undetermined. Surprisingly, our transcriptome analysis has also highlighted a significant enrichment in upregulated genes associated with the synapse suggesting that *FMR1* KO cells, aside from presenting with an impairment in cortical differentiation, may be committed to other neuronal subtypes. We have then validated, in *FMR1* KO neurons, a significant increase in TH expression, required in dopaminergic neurons. Interestingly, in HPRT KO iPSC derived neurons-a cellular model of Lesh-Nyhan disease - the dopaminergic differentiation is impaired due to an inhibition of the mTOR pathway [42]. Reversely, the mTOR pathway was shown to be overactivated in *Fmr1* KO mice due to a lack of FMRP-dependent translational repression [43]. Furthermore, impairments in dopamine signaling have been reported in an *FMR1* KO mouse model [44]. Several experiments need to be done to determine how the absence of FMRP protein could concomitantly affect the cortical differentiation and dopamine signaling which is sensitive to the mTOR pathway [45].

### Neuronal differentiation and electrical activity impairment

Altered neuronal development has been reported in FXS model; and differing findings showing both hypoactivity and hyperactivity have been reported [46, 47]. We observed that cortical neurons from *FMR1* KO cells and FXS patient cells display altered neuronal morphology with defective neurite growth. This impaired neuronal development is associated with hypoactivity. Interestingly, Gildin et al showed that *FMR1* KO induced neurons generated with the overexpression of NGN1 displayed hyperexcitable but less synchronous networks at later stages of development [48]. This highlights how different methodologies to generate mature neurons might influence their phenotype, especially when the progenitor stage is suppressed from the protocol, and we observe important alterations already at this stage.

Even though we observed impaired neuronal activity, we found increased expression of many presynaptic genes in our *FMR1* KO model. Amongst them was the *SLC17A7* gene, which encodes the vesicular transporter of glutamate, and which has also been described as an FMRP target [49]. We also observed a reduction in the GABAergic neuron population in *FMR1* KO. This was consistent with another study that reported an increase in the presynaptic protein VGLUT-1 in iPSC *FMR1* KO cells that also presented with altered synaptic development, but no alteration in the GABAergic marker GAD67 [50]. Thus, the increased expression observed in our model is highly likely due to the lack of FMRP repression in the *FMR1* KO.

### Impaired transcriptome and cellular activity: common traits between FXS and other neurodevelopmental disorders

Our transcriptomic analysis has revealed a decrease in the expression of genes involved in synapse functioning, consistent with many other studies with iPSC-derived cells from patients with idiopathic autism. Furthermore, we have identified, in the list of downregulated genes, significant enrichments in FMRP targets and autism-related genes. We also observed in our *FMR1* KO and FXS patent iPSC-derived neurons decreased activity blocked by TTX and a decrease in the expression of NMDA receptor subunit *GRIN2B*, a gene previously associated with autism [51]. In a previous study, we had shown that *GRIN2B* mutation affects the NMDA response in iPSC-derived cells. Moreover, a chronic APV treatment (an NMDA antagonist) was shown to affect the neuronal differentiation of glutamatergic neurons, similar to a mutation of the *GRIN2B* gene [17]. This demonstrates that at early developmental stages, although the synapses are not formed yet, AMPA and NMDA-dependent activities are impaired in fragile X syndrome and other forms of autism spectrum disorders, and are needed for proper neuronal differentiation. Both signaling pathways could constitute therapeutic targets that must be further assessed by drug screening.

### Advantages of the iPSC model and Perspectives

iPSC-derived models have proven highly useful for investigating the cellular and molecular deregulation associated with neurodevelopmental disorders (**Figure 9**). In our study we successfully analyzed the cellular activity and expression profile of iPSC-derived NPCs and neurons from *FMR1* KO and FXS patient lines, focusing on cortical differentiation. Our study also shows that the lack of FMRP expression affects cortical differentiation during the initial stages of development. Our study also confirmed that the *FMR1* KO, which is largely used as a model of Fragile X syndrome, could pinpoint common deregulations and patterns shared among different autism spectrum disorders through the deregulation of autism-related genes and FMRP targets. However, our study also demonstrated how the absence of FMRP protein could affect the developmental program of cells from the CNS (forebrain specification and increase of genes related to the somatodendritic compartment) raising many unanswered questions. To better understand how the silencing of the *FMR1* gene affects neuronal differentiation and brain regional specification, as well as the development of neuronal and non-neuronal cells within the central nervous system, we would benefit from generating cortical and mesencephalic organoids from *FMR1* iPSC KO cells as more complex models allow the study of cellular and molecular abnormalities [53]. An advantage of brain organoids is their capacity to recapitulate the multilayer organization of the brain [54, 55]. This approach provides the means to study many of the different cellular populations that drive brain development and promote maturation and survival, as well as their connectivity and electrophysiological activity.

In the current study, by combining the iPSC and genome editing technologies we were capable of investigating, the consequences of the *FMR1* absence on the activities and expression profiles of neural progenitors and early differentiating neurons. We have also found that *FMR1* knock-out recapitulates changes in the activities and expression profiles observed in iPSC-derived cells from Fragile X patient. Then, *FMR1*-lacking 3D models would help to further explore how the transcriptional dysregulations and impaired neuronal activities affect (i) the developmental trajectories of neuronal and non-neuronal cells, and (ii) the organization of neural networks in the developing brain.

## Acknowledgments

ABIF - McGill microscopy platform, T.M.D. received funding to support this project through the Canada First Research Excellence Fund, awarded through the Healthy Brains, Healthy Lives initiative at McGill University, the Alain and Sandra Bouchard Foundation, the Chamandy Foundation, and the Djavad Mowafaghian Foundation. Figure 9 was created with BioRender.com.

## Supplementary Information

### Supplementary Tables

**Supplementary Table 1.**
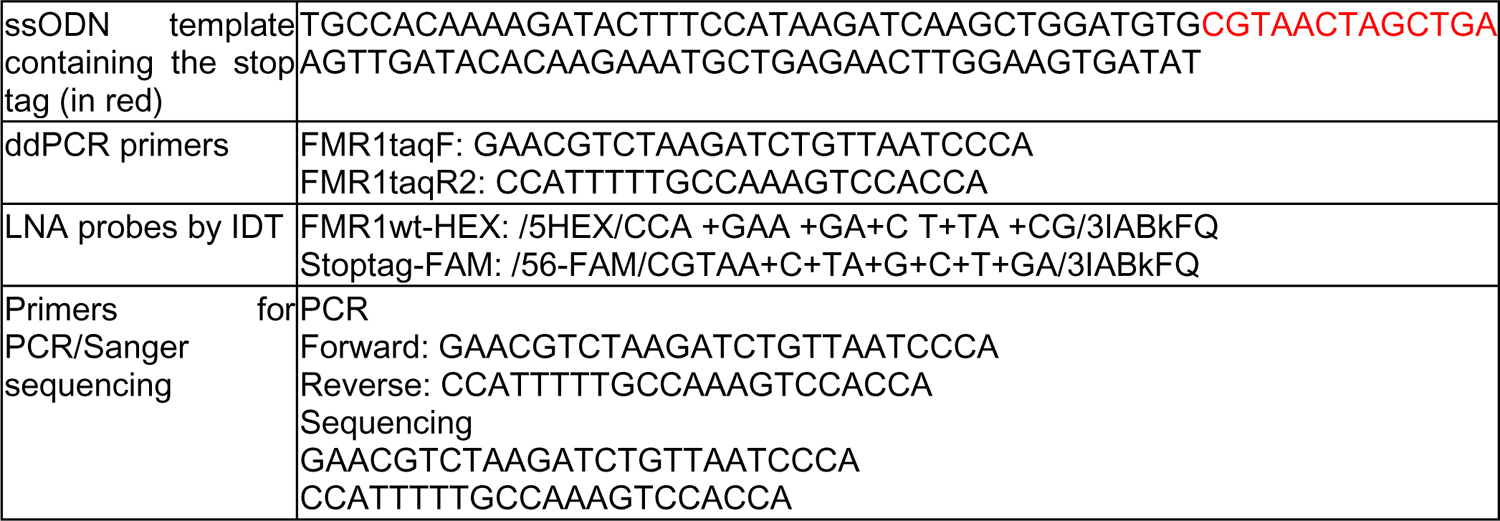
Nucleotide sequences used for Cripr editing, digital PCR, and Sanger sequencing.

**Supplementary Table 2.**
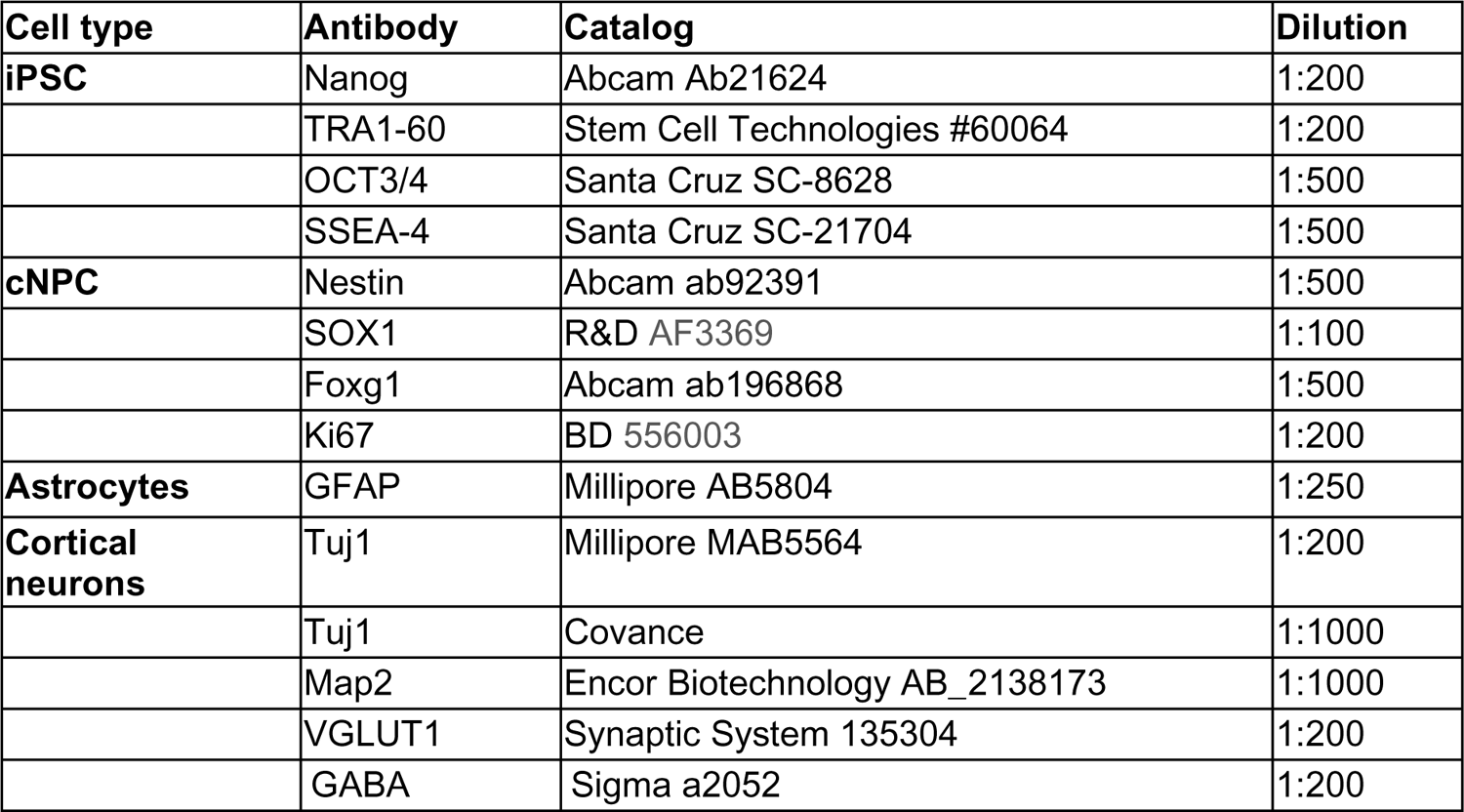
List of antibodies.

| Cell type | Antibody | Catalog | Dilution |
| --- | --- | --- | --- |
| **iPSC** | Nanog | Abcam Ab21624 | 1:200 |
|  | TRA1-60 | Stem Cell Technologies #60064 | 1:200 |
|  | OCT3/4 | Santa Cruz SC-8628 | 1:500 |
|  | SSEA-4 | Santa Cruz SC-21704 | 1:500 |
| **cNPC** | Nestin | Abcam ab92391 | 1:500 |
|  | SOX1 | R&D AF3369 | 1:100 |
|  | Foxg1 | Abcam ab196868 | 1:500 |
|  | Ki67 | BD 556003 | 1:200 |
| **Astrocytes** | GFAP | Millipore AB5804 | 1:250 |
| **Cortical neurons** | Tuj1 | Millipore MAB5564 | 1:200 |
|  | Tuj1 | Covance | 1:1000 |
|  | Map2 | Encor Biotechnology AB_2138173 | 1:1000 |
|  | VGLUT1 | Synaptic System 135304 | 1:200 |
|  | GABA | Sigma a2052 | 1:200 |

**Supplementary Table 3.**
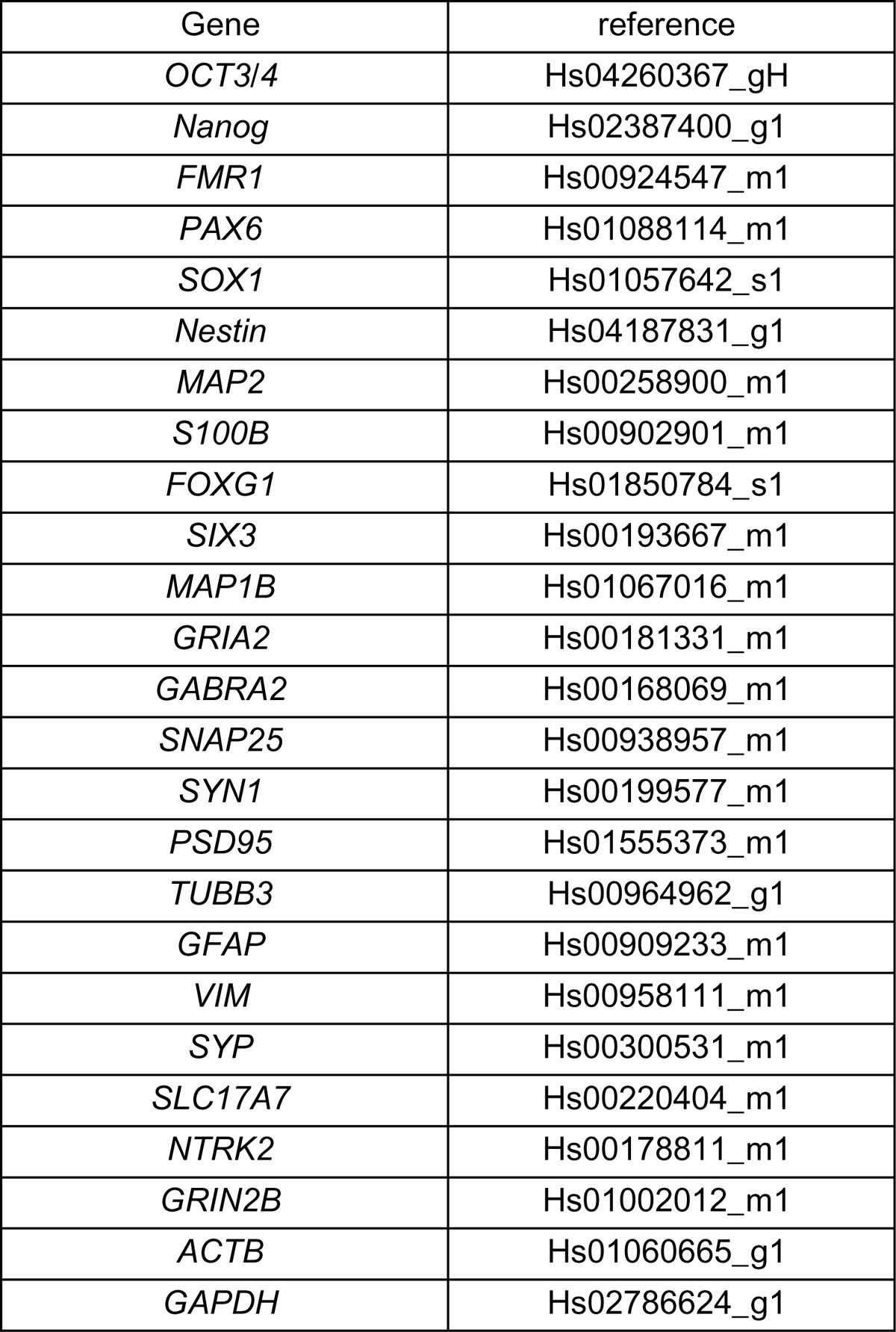
List of TaqMan probes and primer sets (Applied Biosystems).

## Supplementary Methods

### Calcium Imaging

#### Cell culture preparation and Fluo4 incubation

Cells were seeded in 35 mm MatTek Dishes (MatTek corporation) in progenitor medium. They were switched to differentiation medium 24 hours later. On the day of the acquisition, the Fluo4 calcium indicator (Thermo Fisher Scientific, Waltham, MA, USA) was incubated for 30 min at a final concentration of 1 µM. Cells were then washed twice for 5 min with the differentiation medium before acquisition.

#### Acquisition and Drug application

The acquisition was performed using a Zeiss Axio Observer z1 microscope assisted by the Zen 2 software. Pictures were collected at every 400 ms for 5 minutes, with a correction for defined focus every 30 pictures. In photo number 120, NMDA was applied at a final concentration of 2µM.

#### Movies treatment, Data processing, and Statistical analyses

The acquisitions were treated using Fiji/Image J software. Threshold was set up to get rid of the empty space. To assess the NMDA response, we performed an automatic segmentation of the cells to define the regions of interest. Data was collected through the particle analysis module. The ROI manager’s multiple measurement tool was used to measure the average pixel values of each region of interest in each frame of the time stack.

Once the data was extracted from the time stack, the background was subtracted from every single region of interest at every time point. Signal variation is expressed as “ΔF/F0”; F0 being the minimal intensity signal for a given region of interest after background subtraction and ΔF being the difference between an intensity signal at a given timepoint and F0. The amplitude of ΔF/F0 variations following NMDA applications was monitored in the regions of interest and averaged to compare responses between cell lines. Statistical comparisons between the differentiation stages were performed through SPSS 20, using an ANOVA followed by a post-hoc T-Test.

### Supplementary results

#### Common deregulations between *FMR1* KO NPCs and *FMR1* KO neurons

The analysis of our RNA sequencing data shows that a similar number of genes were differentially expressed at the NPC and differentiating neuron stages (**Supplementary** Figure 2). Those genes include genes associated with synapse and neuronal formation, autism-associated genes, and FMRP targets. We then wondered how common the deregulated genes in *FMR1* KO cell lines at the NPC and neuronal stages are. Comparing the two gene lists (Figure 6A **and** Figure 6B), we found a set of 536 genes (40% of the DEGs) that were deregulated at both the NPC and neuronal stages (**Supplementary** Figure 2). We further examined the fold changes in the list of commonly deregulated genes. We found that 88% of them were either up or downregulated at both stages (**Supplementary** Figure 2). Genes that were found upregulated in *FMR1* KO NPCs and neurons are involved in brain development, neurogenesis, and forebrain specification (**Supplementary** Figure 2). These genes include *SIX3* and *FOXG1* which code for transcription factors whose mutations are respectively responsible for holoprosencephaly/schizencephaly [1], Rett-like, and FOXG1 [2] syndromes. We have actually validated by Q-PCR the increased expression of *SIX3* and *FOXG1* in *FMR1* KO NPCs and Neurons (F*_SIX3_* _Genotype_=1145; df=1; p<0.0001; t_NPCs_=18.27; p<0.001; t_Neurons_=29.60; p<0.001; F*_FOXG1_* _Genotype_=264.4; df=1; p<0.0001; t_NPCs_=7.491; p<0.001;t_Neurons_=15.50;p<0.001). Genes downregulated in the *FMR1* KO NPCs and neurons were significantly enriched in synaptic function, axon, neuronal projections, and DNA binding transcription factor (**Supplementary** Figure 2). These functional enrichments observed from common deregulations in *FMR1* KO NPCs and neurons suggest that the regional specification of *FMR1* KO cells and their neuronal differentiation are impaired, leaving the cells in a progenitor-like state. We have also assessed these gene lists for enrichment in ASD-related genes and FMRP targets. Similarly, to what was observed in Figure 6, we found significant enrichment in ASD-related genes (39/262; p < 9.309e-05) (**Supplementary** Figure 2E) and in FMRP targets (73/262; p < 2.526e-07) (**Supplementary** Figure 2G) among the list of genes that are commonly downregulated in *FMR1* KO NPCs and neurons.

## Legends of supplementary figures

**Supplementary Figure 1:**
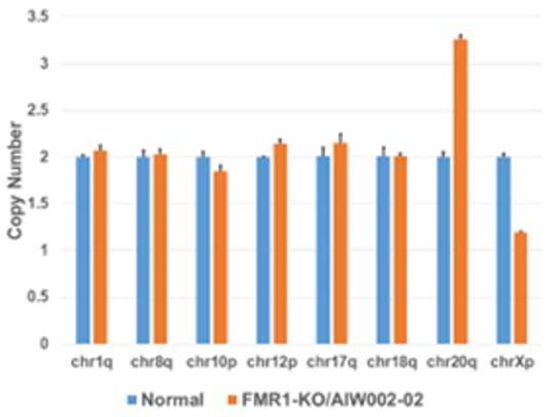
Genomic stability testing of the *FMR1* KO iPSC line. qPCR-based chromosomal abnormality analysis in the eight common critical areas within chr1q, chr8q, chr10p, chr12p, chr17q, chr18q, chr20q, and chrXp showed amplification in chr20q only.

**Supplementary Figure 2:**
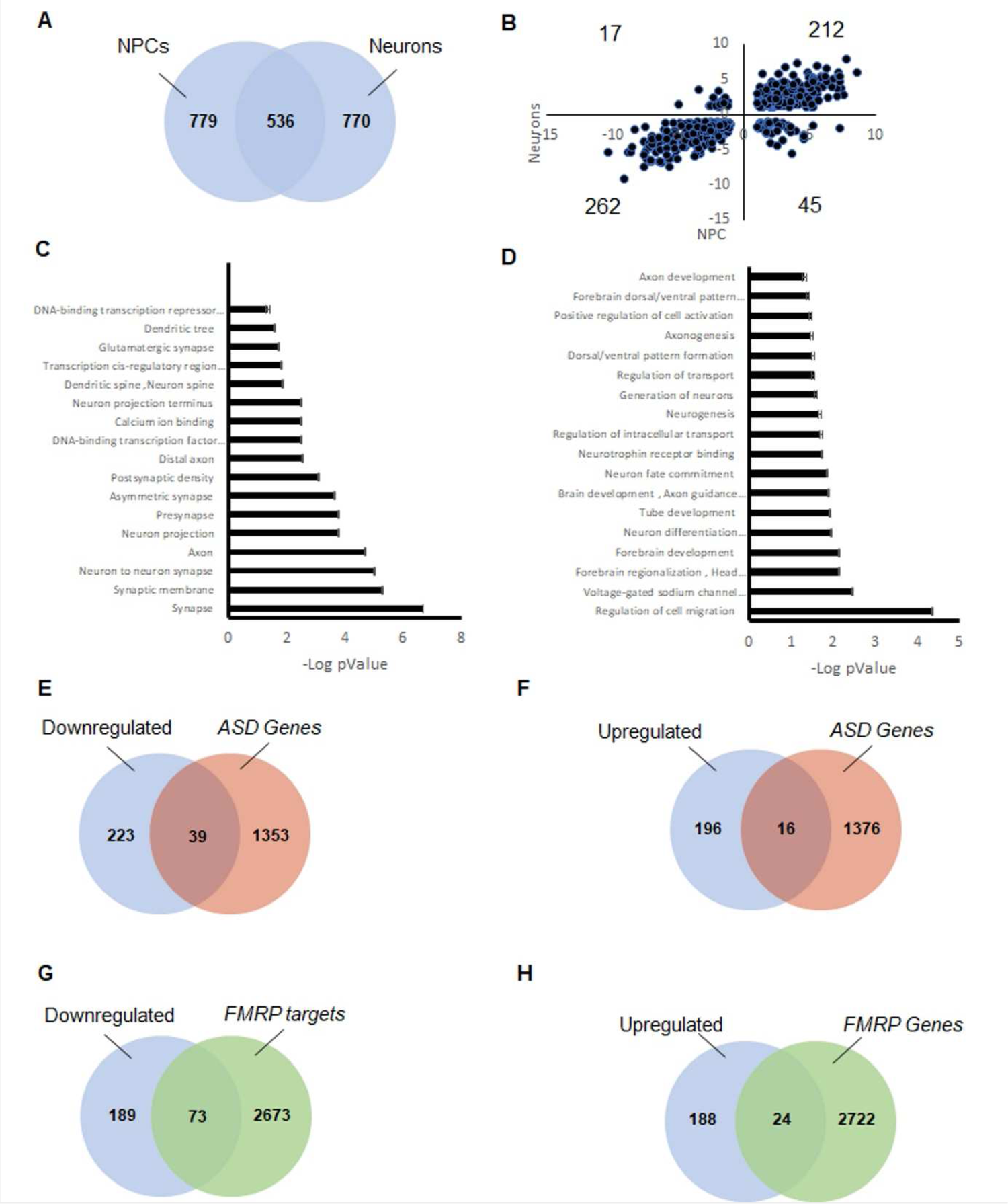
Common transcriptional deregulations between *FMR1*-KO NPCs and Neurons. A) Venn diagram of the differentially expressed genes in *FMR1* KO NPCs and *FMR1* KO neurons. B) Fold change distribution for the common set of genes differentially expressed in *FMR1* KO NPCs and Neurons. (C) GO terms defining the significant functional enrichment for the genes that are downregulated in *FMR1* KO NPCs and *FMR1* KO neurons. (D) GO terms defining the significant functional enrichment for the genes that are upregulated in *FMR1* KO NPCs and *FMR1* KO neurons. (E) Overlap between the genes significantly downregulated in *FMR1* KO NPCs and neurons, and the list of autism risk genes analyzed. (F) Overlap between the genes significantly upregulated in *FMR1* KO NPCs and neurons, and the list of autism risk genes. (G) Overlap between the genes significantly downregulated in *FMR1* KO NPCs and neurons, and the list of FMRP targets analyzed. (H) Overlap between the genes significantly upregulated in *FMR1* KO NPCs and neurons, and the list of FMRP targets analyzed.

**Supplementary Figure 3:**
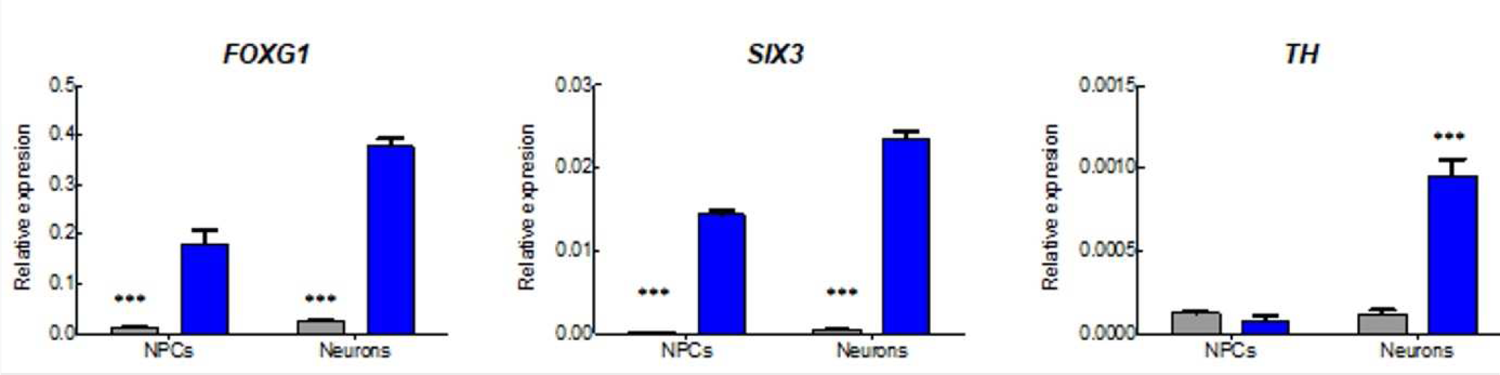
Validation of differential gene expression by Real-Time PCR. Normalized expression levels of *FOXG1*, *SIX3*, and *TH* mRNA in *FMR1* KO NPCs and neurons compared to control cells.

**Supplementary Figure 4:**
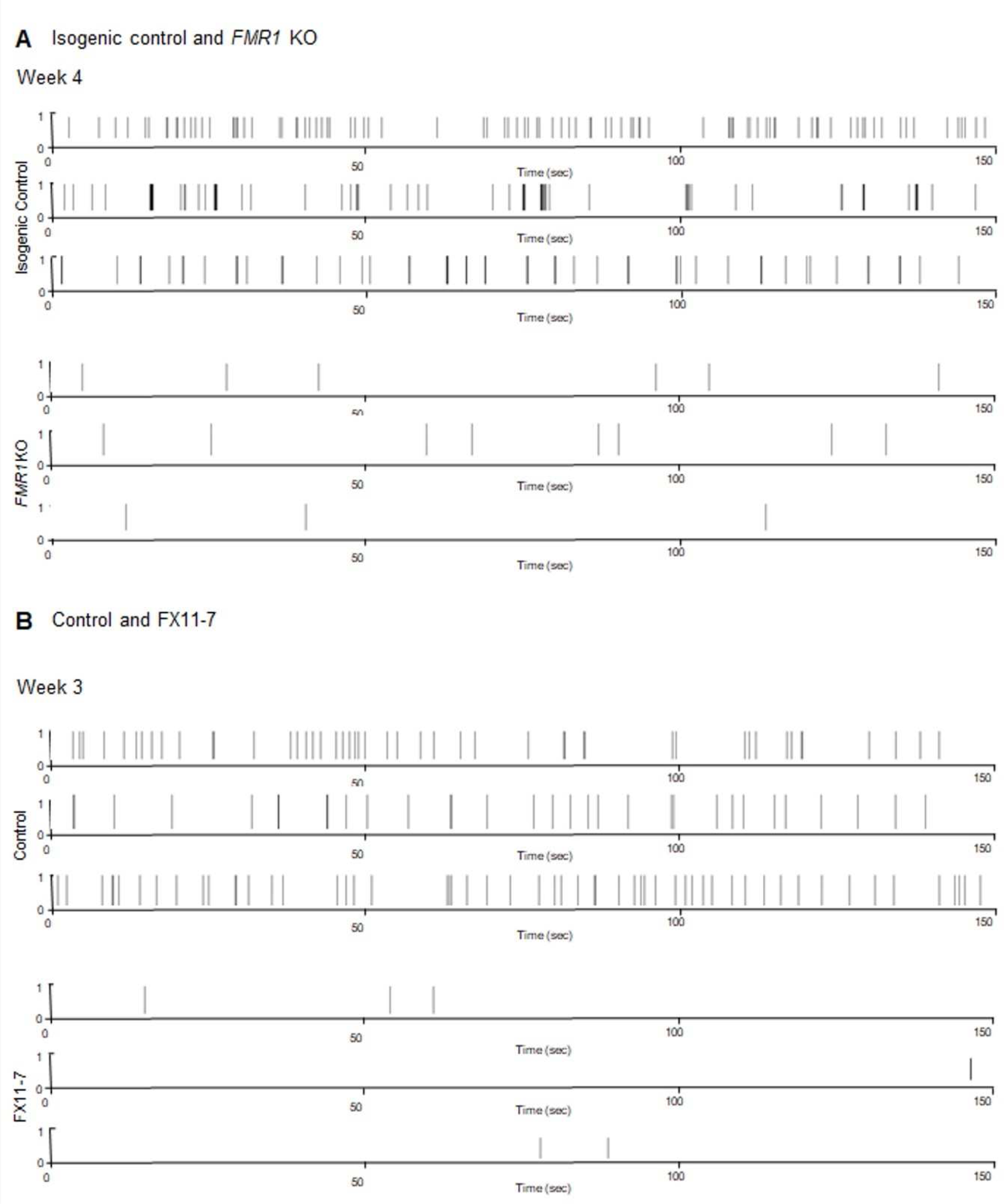
Validation of differential gene expression by Real-Time PCR. MEA raster plots from MEA recordings showing. Changes in electrical activity in week 3 to week 4 in cortical neurons from (A) Isogenic control and *FMR1* KO, and (B) Control and FX11-7.

**Supplementary Figure 5:**
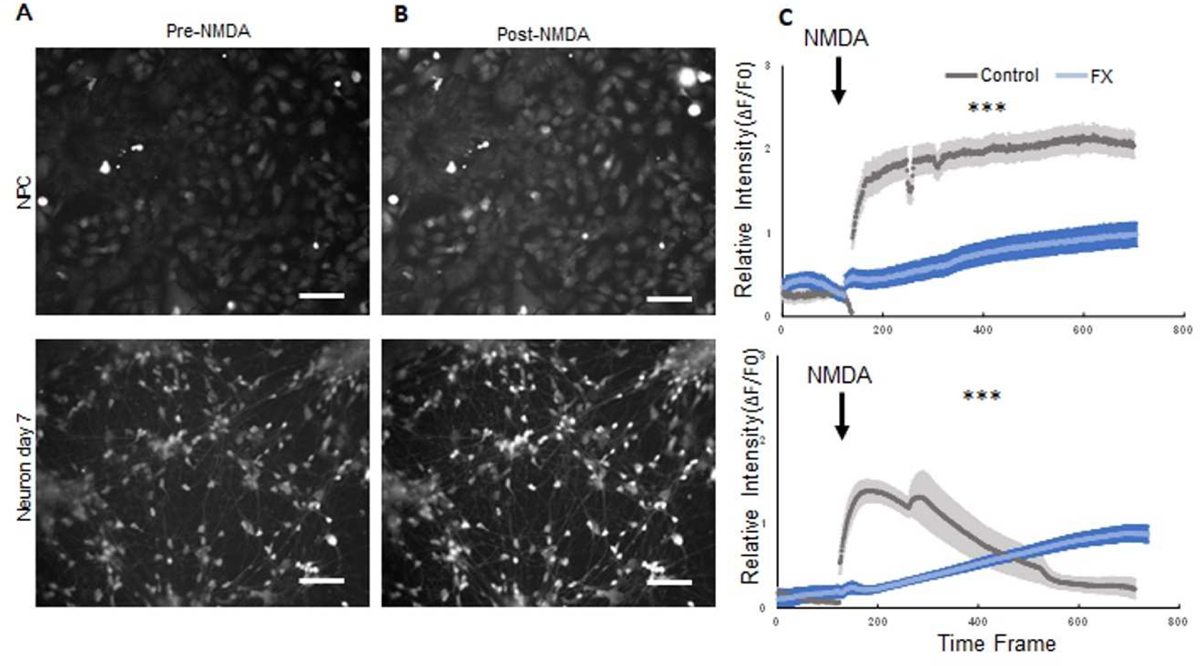
Differential NMDA response between control and FXS IPSC-derived NPCs and Neurons. (A-B) Manual segmentation of the regions of interest before and after NMDA application. (C) Graphical representation of the averaged relative activity expressed as ΔF/F0 during the time of the acquisition. Scale bar: 10µM

## References

1. Abrahams, B. S.; Geschwind, D. H., Advances in autism genetics: on the threshold of a new neurobiology. Nat Rev Genet 2008, 9, (5), 341–55.

2. Verkerk, A. J.; Pieretti, M.; Sutcliffe, J. S.; Fu, Y. H.; Kuhl, D. P.; Pizzuti, A.; Reiner, O.; Richards, S.; Victoria, M. F.; Zhang, F. P.;, et al., Identification of a gene (FMR-1) containing a CGG repeat coincident with a breakpoint cluster region exhibiting length variation in fragile X syndrome. Cell 1991, 65, (5), 905–14.

3. Davis, J. K.; Broadie, K., Multifarious Functions of the Fragile X Mental Retardation Protein. Trends Genet 2017, 33, (10), 703–714.

4. Richter, J. D.; Zhao, X., The molecular biology of FMRP: new insights into fragile X syndrome. Nat Rev Neurosci 2021, 22, (4), 209–222.

5. Khandjian, E. W.; Robert, C.; Davidovic, L., FMRP, a multifunctional RNA-binding protein in quest of a new identity. Front Genet 2022, 13, 976480.

6. Wang, L. W.; Berry-Kravis, E.; Hagerman, R. J., Fragile X: leading the way for targeted treatments in autism. Neurotherapeutics 2010, 7, (3), 264–74.

7. Schaefer, G. B.; Mendelsohn, N. J., Genetics evaluation for the etiologic diagnosis of autism spectrum disorders. Genet Med 2008, 10, (1), 4–12.

8. Penagarikano, O.; Mulle, J. G.; Warren, S. T., The pathophysiology of fragile x syndrome. Annu Rev Genomics Hum Genet 2007, 8, 109–29.

9. Maussion, G.; Rocha, C.; Bernard, G.; Beitel, L. K.; Durcan, T. M., Patient-Derived Stem Cells, Another in vitro Model, or the Missing Link Toward Novel Therapies for Autism Spectrum Disorders? Front Pediatr 2019, 7, 225.

10. Maussion, G.; Rocha, C.; Pimentel, L.; Beitel, L. K.; Durcan, T. M., Chapter 3 - Human induced pluripotent stem cell-based studies; a new route toward modeling autism spectrum disorders. In iPSCs for Modeling Central Nervous System Disorders, Birbrair, A., Ed. Academic Press: 2021; Vol. 6, pp 37–81.

11. Takahashi, K.; Tanabe, K.; Ohnuki, M.; Narita, M.; Ichisaka, T.; Tomoda, K.; Yamanaka, S., Induction of pluripotent stem cells from adult human fibroblasts by defined factors. Cell 2007, 131, (5), 861–72.

12. Ardhanareeswaran, K.; Mariani, J.; Coppola, G.; Abyzov, A.; Vaccarino, F. M., Human induced pluripotent stem cells for modelling neurodevelopmental disorders. Nat Rev Neurol 2017, 13, (5), 265–278.

13. Shi, Y.; Inoue, H.; Wu, J. C.; Yamanaka, S., Induced pluripotent stem cell technology: a decade of progress. Nat Rev Drug Discov 2017, 16, (2), 115–130.

14. Li, Y.; Wang, R.; Qiao, N.; Peng, G.; Zhang, K.; Tang, K.; Han, J. J.; Jing, N., Transcriptome analysis reveals determinant stages controlling human embryonic stem cell commitment to neuronal cells. J Biol Chem 2017, 292, (48), 19590–19604.

15. Chen, E. S.; Gigek, C. O.; Rosenfeld, J. A.; Diallo, A. B.; Maussion, G.; Chen, G. G.; Vaillancourt, K.; Lopez, J. P.; Crapper, L.; Poujol, R.; Shaffer, L. G.; Bourque, G.; Ernst, C., Molecular convergence of neurodevelopmental disorders. Am J Hum Genet 2014, 95, (5), 490–508.

16. Gigek, C. O.; Chen, E. S.; Ota, V. K.; Maussion, G.; Peng, H.; Vaillancourt, K.; Diallo, A. B.; Lopez, J. P.; Crapper, L.; Vasuta, C.; Chen, G. G.; Ernst, C., A molecular model for neurodevelopmental disorders. Transl Psychiatry 2015, 5, (5), e565.

17. Bell, S.; Maussion, G.; Jefri, M.; Peng, H.; Theroux, J. F.; Silveira, H.; Soubannier, V.; Wu, H.; Hu, P.; Galat, E.; Torres-Platas, S. G.; Boudreau-Pinsonneault, C.; O’Leary, L. A.; Galat, V.; Turecki, G.; Durcan, T. M.; Fon, E. A.; Mechawar, N.; Ernst, C., Disruption of GRIN2B Impairs Differentiation in Human Neurons. Stem Cell Reports 2018, 11, (1), 183–196.

18. Tang, B.; Wang, T.; Wan, H.; Han, L.; Qin, X.; Zhang, Y.; Wang, J.; Yu, C.; Berton, F.; Francesconi, W.; Yates, J. R 3rd.,; Vanderklish, P. W.; Liao, L., Fmr1 deficiency promotes age-dependent alterations in the cortical synaptic proteome. Proc Natl Acad Sci USA 2015, 112, (34), E4697–706.

19. Mor-Shaked, H.; Eiges, R., Modeling Fragile X Syndrome Using Human Pluripotent Stem Cells. Genes (Basel) 2016, 7, (10).

20. Volpato, V.; Webber, C., Addressing variability in iPSC-derived models of human disease: guidelines to promote reproducibility. Dis Model Mech 2020, 13, (1).

21. Autar, K.; Guo, X.; Rumsey, J. W.; Long, C. J.; Akanda, N.; Jackson, M.; Narasimhan, N. S.; Caneus, J.; Morgan, D.; Hickman, J. J., A functional hiPSC-cortical neuron differentiation and maturation model and its application to neurological disorders. Stem Cell Reports 2022, 17, (1), 96–109.

22. Chen, C. X.; Abdian, N.; Maussion, G.; Thomas, R. A.; Demirova, I.; Cai, E.; Tabatabaei, M.; Beitel, L. K.; Karamchandani, J.; Fon, E. A.; Durcan, T. M., A Multistep Workflow to Evaluate Newly Generated iPSCs and Their Ability to Generate Different Cell Types. Methods Protoc 2021, 4, (3).

23. Bell, S.; Peng, H.; Crapper, L.; Kolobova, I.; Maussion, G.; Vasuta, C.; Yerko, V.; Wong, T. P.; Ernst, C., A Rapid Pipeline to Model Rare Neurodevelopmental Disorders with Simultaneous CRISPR/Cas9 Gene Editing. Stem Cells Transl Med 2017, 6, (3), 886–896.

24. Pellegrini, R., Edit single bases with Benchling! 2016.

25. Deneault, E.; Chaineau, M.; Nicouleau, M.; Castellanos Montiel, M. J.; Franco Flores, A. K.; Haghi, G.; Chen, C. X.; Abdian, N.; Shlaifer, I.; Beitel, L. K.; Durcan, T. M., A streamlined CRISPR workflow to introduce mutations and generate isogenic iPSCs for modeling amyotrophic lateral sclerosis. Methods 2022, 203, 297–310.

26. Nicouleau, M.; Pimentel, L.; Shlaifer, I.; Durcan, T. M., Generation of Knockout Cell Lines Using CRISPR-Cas9 and ddPCR Technology. 2020.

27. Nicouleau, M.; Durcan, T. M., DNA sequencing with the SeqStudio. 2020.

28. Maussion, G.; Thomas, R. A.; Demirova, I.; Gu, G.; Cai, E.; Chen, C. X.; Abdian, N.; Strauss, T. J. P.; Kelai, S.; Nauleau-Javaudin, A.; Beitel, L. K.; Ramoz, N.; Gorwood, P.; Durcan, T. M., Auto-qPCR; a python-based web app for automated and reproducible analysis of qPCR data. Sci Rep 2021, 11, (1), 21293.

29. Li, H., Minimap2: pairwise alignment for nucleotide sequences. Bioinformatics 2018, 34, (18), 3094–3100.

30. Liao, Y.; Smyth, G. K.; Shi, W., featureCounts: an efficient general purpose program for assigning sequence reads to genomic features. Bioinformatics 2014, 30, (7), 923–30.

31. Love, M. I.; Huber, W.; Anders, S., Moderated estimation of fold change and dispersion for RNA-seq data with DESeq2. Genome Biol 2014, 15, (12), 550.

32. Benjamini, Y.; Hochberg, Y., Controlling the False Discovery Rate: A Practical and Powerful Approach to Multiple Testing. Journal of the Royal Statistical Society: Series B (Methodological) 1995, 57, (1).

33. Ge, S. X.; Jung, D.; Yao, R., ShinyGO: a graphical gene-set enrichment tool for animals and plants. Bioinformatics 2020, 36, (8), 2628–2629.

34. Basu, S. N.; Kollu, R.; Banerjee-Basu, S., AutDB: a gene reference resource for autism research. Nucleic Acids Res 2009, 37, (Database issue), D832–6.

35. Tran, S. S.; Jun, H. I.; Bahn, J. H.; Azghadi, A.; Ramaswami, G.; Van Nostrand, E. L.; Nguyen, T. B.; Hsiao, Y. E.; Lee, C.; Pratt, G. A.; Martinez-Cerdeno, V.; Hagerman, R. J.; Yeo, G. W.; Geschwind, D. H.; Xiao, X., Widespread RNA editing dysregulation in brains from autistic individuals. Nat Neurosci 2019, 22, (1), 25–36.

36. Bardy, C.; van den Hurk, M.; Eames, T.; Marchand, C.; Hernandez, R. V.; Kellogg, M.; Gorris, M.; Galet, B.; Palomares, V.; Brown, J.; Bang, A. G.; Mertens, J.; Bohnke, L.; Boyer, L.; Simon, S.; Gage, F. H., Neuronal medium that supports basic synaptic functions and activity of human neurons in vitro. Proc Natl Acad Sci U S A 2015, 112, (20), E2725–34.

37. Hyvarinen, T.; Hyysalo, A.; Kapucu, F. E.; Aarnos, L.; Vinogradov, A.; Eglen, S. J.; Yla-Outinen, L.; Narkilahti, S., Functional characterization of human pluripotent stem cell-derived cortical networks differentiated on laminin-521 substrate: comparison to rat cortical cultures. Sci Rep 2019, 9, (1), 17125.

38. Harrell, E. R.; Pimentel, D.; Miesenbock, G., Changes in Presynaptic Gene Expression during Homeostatic Compensation at a Central Synapse. J Neurosci 2021, 41, (14), 3054–3067.

39. Kasteel, E. E.; Westerink, R. H., Comparison of the acute inhibitory effects of Tetrodotoxin (TTX) in rat and human neuronal networks for risk assessment purposes. Toxicol Lett 2017, 270, 12–16.

40. Weisz, E. D.; Monyak, R. E.; Jongens, T. A., Deciphering discord: How Drosophila research has enhanced our understanding of the importance of FMRP in different spatial and temporal contexts. Exp Neurol 2015, 274, (Pt A), 14–24.

41. Raj, N.; McEachin, Z. T.; Harousseau, W.; Zhou, Y.; Zhang, F.; Merritt-Garza, M. E.; Taliaferro, J. M.; Kalinowska, M.; Marro, S. G.; Hales, C. M.; Berry-Kravis, E.; Wolf-Ochoa, M. W.; Martinez-Cerdeno, V.; Wernig, M.; Chen, L.; Klann, E.; Warren, S. T.; Jin, P.; Wen, Z.; Bassell, G. J., Cell-type-specific profiling of human cellular models of fragile X syndrome reveal PI3K-dependent defects in translation and neurogenesis. Cell Rep 2021, 35, (2), 108991.

42. Bell, S.; McCarty, V.; Peng, H.; Jefri, M.; Hettige, N.; Antonyan, L.; Crapper, L.; O’Leary, L. A.; Zhang, X.; Zhang, Y.; Wu, H.; Sutcliffe, D.; Kolobova, I.; Rosenberger, T. A.; Moquin, L.; Gratton, A.; Popic, J.; Gantois, I.; Stumpf, P. S.; Schuppert, A. A.; Mechawar, N.; Sonenberg, N.; Tremblay, M. L.; Jinnah, H. A.; Ernst, C., Lesch-Nyhan disease causes impaired energy metabolism and reduced developmental potential in midbrain dopaminergic cells. Stem Cell Reports 2021, 16, (7), 1749–1762.

43. Sharma, A.; Hoeffer, C. A.; Takayasu, Y.; Miyawaki, T.; McBride, S. M.; Klann, E.; Zukin, R. S., Dysregulation of mTOR signaling in fragile X syndrome. J Neurosci 2010, 30, (2), 694–702.

44. Chao, O. Y.; Pathak, S. S.; Zhang, H.; Dunaway, N.; Li, J. S.; Mattern, C.; Nikolaus, S.; Huston, J. P.; Yang, Y. M., Altered dopaminergic pathways and therapeutic effects of intranasal dopamine in two distinct mouse models of autism. Mol Brain 2020, 13, (1), 111.

45. Kosillo, P.; Ahmed, K. M.; Aisenberg, E. E.; Karalis, V.; Roberts, B. M.; Cragg, S. J.; Bateup, H. S., Dopamine neuron morphology and output are differentially controlled by mTORC1 and mTORC2. Elife 2022, 11.

46. Doers, M. E.; Musser, M. T.; Nichol, R.; Berndt, E. R.; Baker, M.; Gomez, T. M.; Zhang, S. C.; Abbeduto, L.; Bhattacharyya, A., iPSC-derived forebrain neurons from FXS individuals show defects in initial neurite outgrowth. Stem Cells Dev 2014, 23, (15), 1777–87.

47. Zhang, Z.; Marro, S. G.; Zhang, Y.; Arendt, K. L.; Patzke, C.; Zhou, B.; Fair, T.; Yang, N.; Sudhof, T. C.; Wernig, M.; Chen, L., The fragile X mutation impairs homeostatic plasticity in human neurons by blocking synaptic retinoic acid signaling. Sci Transl Med 2018, 10, (452).

48. Gildin, L.; Rauti, R.; Vardi, O.; Kuznitsov-Yanovsky, L.; Maoz, B. M.; Segal, M.; Ben-Yosef, D., Impaired Functional Connectivity Underlies Fragile X Syndrome. Int J Mol Sci 2022, 23, (4).

49. Tao, J.; Wu, H.; Coronado, A. A.; de Laittre, E.; Osterweil, E. K.; Zhang, Y.; Bear, M. F., Negative Allosteric Modulation of mGluR5 Partially Corrects Pathophysiology in a Mouse Model of Rett Syndrome. J Neurosci 2016, 36, (47), 11946–11958.

50. Brighi, C.; Salaris, F.; Soloperto, A.; Cordella, F.; Ghirga, S.; de Turris, V.; Rosito, M.; Porceddu, P. F.; D’Antoni, C.; Reggiani, A.; Rosa, A.; Di Angelantonio, S., Novel fragile X syndrome 2D and 3D brain models based on human isogenic FMRP-KO iPSCs. Cell Death Dis 2021, 12, (5), 498.

51. Pan, Y.; Chen, J.; Guo, H.; Ou, J.; Peng, Y.; Liu, Q.; Shen, Y.; Shi, L.; Liu, Y.; Xiong, Z.; Zhu, T.; Luo, S.; Hu, Z.; Zhao, J.; Xia, K., Association of genetic variants of GRIN2B with autism. Sci Rep 2015, 5, 8296.

52. O’Roak, B. J.; Vives, L.; Fu, W.; Egertson, J. D.; Stanaway, I. B.; Phelps, I. G.; Carvill, G.; Kumar, A.; Lee, C.; Ankenman, K.; Munson, J.; Hiatt, J. B.; Turner, E. H.; Levy, R.; O’Day, D. R.; Krumm, N.; Coe, B. P.; Martin, B. K.; Borenstein, E.; Nickerson, D. A.; Mefford, H. C.; Doherty, D.; Akey, J. M.; Bernier, R.; Eichler, E. E.; Shendure, J., Multiplex targeted sequencing identifies recurrently mutated genes in autism spectrum disorders. Science 2012, 338, (6114), 1619–22.

53. Kang, Y.; Zhou, Y.; Li, Y.; Han, Y.; Xu, J.; Niu, W.; Li, Z.; Liu, S.; Feng, H.; Huang, W.; Duan, R.; Xu, T.; Raj, N.; Zhang, F.; Dou, J.; Xu, C.; Wu, H.; Bassell, G. J.; Warren, S. T.; Allen, E. G.; Jin, P.; Wen, Z., A human forebrain organoid model of fragile X syndrome exhibits altered neurogenesis and highlights new treatment strategies. Nat Neurosci 2021, 24, (10), 1377–1391.

54. Lancaster, M. A.; Renner, M.; Martin, C. A.; Wenzel, D.; Bicknell, L. S.; Hurles, M. E.; Homfray, T.; Penninger, J. M.; Jackson, A. P.; Knoblich, J. A., Cerebral organoids model human brain development and microcephaly. Nature 2013, 501, (7467), 373–9.

55. Pasca, A. M.; Sloan, S. A.; Clarke, L. E.; Tian, Y.; Makinson, C. D.; Huber, N.; Kim, C. H.; Park, J. Y.; O’Rourke, N. A.; Nguyen, K. D.; Smith, S. J.; Huguenard, J. R.; Geschwind, D. H.; Barres, B. A.; Pasca, S. P., Functional cortical neurons and astrocytes from human pluripotent stem cells in 3D culture. Nat Methods 2015, 12, (7), 671–8.

## References

1. Hehr, U.; Pineda-Alvarez, D. E.; Uyanik, G.; Hu, P.; Zhou, N.; Hehr, A.; Schell-Apacik, C.; Altus, C.; Daumer-Haas, C.; Meiner, A.; Steuernagel, P.; Roessler, E.; Winkler, J.; Muenke, M., Heterozygous mutations in SIX3 and SHH are associated with schizencephaly and further expand the clinical spectrum of holoprosencephaly. Hum Genet 2010, 127, (5), 555–61.

2. Jacob, F. D.; Ramaswamy, V.; Andersen, J.; Bolduc, F. V., Atypical Rett syndrome with selective FOXG1 deletion detected by comparative genomic hybridization: case report and review of literature. Eur J Hum Genet 2009, 17, (12), 1577–81.

